# G-quadruplexes as pivotal components of *cis*-regulatory elements in the human genome

**DOI:** 10.1101/2024.01.02.573847

**Authors:** Rongxin Zhang, Yuqi Wang, Cheng Wang, Xiao Sun, Jean-Louis Mergny

## Abstract

*Cis*-regulatory elements have the ability to precisely regulate gene expression in cells, and G-quadruplexes (G4s), as non-canonical secondary structures, may potentially play a role in this regulation. However, a lack of systematic studies exists to uncover the connection between these two elements. Here, we comprehensively characterized the associations between G4s and human candidate *cis*-regulatory elements (cCREs) inferred from the Encyclopedia of DNA Elements (ENCODE) data. Our findings demonstrate that G4s are prominently enriched in most types of cCREs, particularly in elements with promoter-like signatures (PLS). Furthermore, we observed that the co-occurrence of CTCF signals with H3K4me3 or H3K27ac signals significantly strengthens the association between cCREs and G4s. This association becomes more pronounced when examining cell type-specific cCREs and G4s. Interestingly, compared to cCREs, genetic variants occurring in G4s, especially within their G-runs, often exhibit higher regulatory potential and deleterious effects. Runs of several consecutive guanines (G-runs) in the vicinity of transcriptional start sites tend to be more constrained in mammalian evolution than cCREs’s G-runs. Besides, the presence of G4s, is often linked to a more favorable local chromatin environment for the activation and execution of regulatory functions of cCREs, potentially attributable to the formation of G4 secondary structures. Finally, we discovered that G4-associated cCREs exhibit widespread activation in a variety of cancers. Altogether, our study suggests that G4s are integral components of human *cis*-regulatory elements, and the G4 primary sequences are associated with the localization of the cCREs, while the G4 structures are linked to the activation of the cCREs. Therefore, we propose to define G4s as pivotal regulatory elements in the human genome.

## Introduction

Eukaryotic gene expression is intricately regulated by *cis*-regulatory elements (CREs) [1, 2]. CREs are sequences located within the non-coding regions of the genome, some of which are positioned near the gene transcription start site (TSS), known as promoters, while others, including enhancers, may be located away from the genes and regulate their expression through spatial and proximal mechanisms, which are facilitated by the three-dimensional chromatin architecture [3]. These *cis*-regulatory elements can interact with transcription factors, acting like switches to precisely control gene expression. Moreover, there are other types of *cis*-regulatory elements, including silencers and insulators [3]. Silencers repress gene expression [4], while insulators isolate two adjacent chromosomal regions to prevent mutual interference in gene regulations [5]. The coordinated operation of *cis*-regulatory elements plays a crucial role in cell differentiation and development, and the utilization of cell-type-specific *cis*-regulatory elements forms the cornerstone of cellular functional diversity.

G-quadruplexes (G4s) are unusual nucleic acid secondary structures, typically formed in guanine-rich regions [6, 7]. The basic unit of G4s is called a G-tetrad or G-quartet, in which four guanines are linked by Hoogsteen hydrogen bonds to form a stable planar - or near-planar - assembly [6, 7]. Multiple G-tetrads can be stacked to create a stable G4 structure. G4 structures often possess canonical primary sequences (*e.g.,* G_x_N_1-7_G_x_N_1-7_G_x_N_1-7_G_x_, where x ≥ 3 and N can be any base). However, the actual situation can be more complex since a number of sequences escaping this consensus may also form stable G4 structures [6, 7]. Therefore, various G4 prediction tools have been developed, including G4Hunter [8], which do not rely entirely on a specific consensus sequence pattern. Computational studies have shown that G4s are not randomly distributed in the human genome; instead, they are concentrated in functional regions such as promoters [9], telomeres [10], and intron-exon borders [11], suggesting that G4s may have biological functions. Furthermore, in recent years, with the widespread application of single-molecule techniques in G4 research and the development of whole genome methods, including G4 ChIP-seq [12] and G4Access [13], more and more evidence indicates that G4s, particularly G4 secondary structures, are not genomic “junk” or accidents but possess important biological functions. Extensive studies have revealed that G4s are widely involved in the regulation of a variety of biological processes, including gene transcription [9], protein translation [14], genome stability [15], telomere regulation [16], and genome replication [17]. However, we currently remain uncertain about whether there is an association between G4s and cCREs, and whether G4s might be the functional component of cCREs. Additionally, it is worth further exploration to determine whether the activation of cCREs is related to the G4 structures or the G4 sequences.

Thanks to the widespread use of sequencing technologies and the ease of storing and sharing epigenetic data, it is now possible to study and annotate the human genome in depth. An international, large-scale study constructed a set of candidate *cis*-regulatory elements (cCREs) for the human and mouse genomes by comprehensively analyzing epigenetic data from the ENCODE (Encyclopedia of DNA Elements) project [18]. This provides us with the opportunity to systematically assess the association between G4s and *cis*-regulatory elements, revealing whether G4s may constitute a key component of *cis*-regulatory elements in the human genome.

In this study, we primarily focused on exploring the importance of G4s in the candidate *cis*-regulatory elements of the human genome. We first determined whether G4 sequences were enriched in various groups of cCREs, as the preferential distribution of G4s often hints at their potential biological functions. Subsequently, we assessed whether variants in G4-associated cCREs were more likely to have regulatory effects as well as more deleterious compared to cCREs themselves using RegulomeDB scores and LINSIGHT scores. Additionally, we interrogated the conservativeness of these cCRE-G4s from an evolutionary perspective based on 241 mammalian species. Furthermore, we investigated whether the presence of G4s, especially the structures, might correlate with a more favorable local chromatin environment for the activation and utilization of cCREs. Finally, we examined whether the activation of cCREs remains closely associated with G4s in multiple cancers. Our research demonstrates that G4s are indeed fundamental components of cCREs, which lays the theoretical foundation for subsequent in-depth studies and interpretations of cCRE functionality.

## Methods

### G-quadruplex sequence prediction

We employed the G4Hunter software [8] for the prediction and identification of all possible G-quadruplex sequences in both the forward and reverse strands of the human genome, with a score threshold set at 1.5 (default value being 1.2, but a threshold of 1.5 ensures the candidate sequences actually form the G4 structures with more than 98% confidence *in vitro*) and a window size of 25 (default value). In this study, only the 22 autosomes and the X chromosome were considered and studied (*i.e.*, the Y-chromosome was excluded due to the unavailability of partial epigenetic datasets). Unless otherwise specified, the assembly version of the reference genome we used was hg38. As a result, a total of 1,435,201 human genome G4 sequences were obtained, comprising 717,551 motifs on the forward strand and 717,650 on the reverse strand, with an average width of 33 base pairs (bp, Supplementary Fig. S1A).

### Endogenous G-quadruplex dataset

The endogenous G4 data for the K562 and HepG2 cell lines were obtained from the Gene Expression Omnibus (GEO) with accession number GSE145090. Unlike the human genomic G4 sequences predicted by G4Hunter, these G4s were captured through G4 ChIP-seq experiments, thereby representing the G4 secondary structures that were formed in specific cellular environments. We utilized high-confidence endogenous G4 data (GSE145090_20180108_K562_async_rep1-3.mult.5of8.bed for K562 cells and GSE145090_HepG2_async_rep1-3.mult.6of9.bed for HepG2 cells) in this study and performed genome assembly version conversion using liftOver software. In this study, when referring to G4s supported by G4 ChIP-seq experiments, it specifically denotes endogenous G4s.

### Candidate *cis*-regulatory elements (cCREs) datasets and annotation

The human candidate *cis*-regulatory elements (cCREs) were retrieved from the SCREEN (Search Candidate *cis*-Regulatory Elements by ENCODE) database (Registry V3), which encompasses five distinct groups: promoter-like (PLS), proximal enhancer-like (pELS), distal enhancer-like (dELS), CTCF-only, and DNase-H3K4me3 (Supplementary Fig. S1B). These various cCREs groups are primarily classified based on genomic signals, including DNase, H3K4me3, H3K27ac, and CTCF, as well as genomic context annotations. In brief, PLS cCREs are characterized by high DNase and high H3K4me3 signals and are located within 200 bp of the nearest transcription start site (TSS). The cCREs with high H3K4me3 signals and low H3K27ac signals, positioned more than 200 bp from the TSS, are considered as DNase-H3K4me3 cCREs. Elements with high H3K27ac signals are regarded as candidate enhancer-like (ELS) cCREs, with their classification also dependent on their distance to the TSS. If the distance is less than 2,000 bp, they will be categorized as pELS (proximal ELS, with low H3K4me3 signal as well). Otherwise, they are designated as dELS (distal ELS). CTCF-only cCREs refer to elements with high CTCF signals but low H3K4me3 and H3K27ac signals. Please note that PLS, ELS, *etc*. may also contain high CTCF signals. Additionally, chromatin accessibility, as indicated by high DNase signals, is a prerequisite for all cCREs elements.

We annotated the presence of G4s in these cCREs (Fig. 1A). For a given cCRE, we will consider it to be a G4-associated cCRE if it overlaps any of the predicted G4 sequences (Fig. 1A). Likewise, this G4 is referred to as a cCRE-associated G4 (or, more briefly in the manuscript, as a cCRE G4, Fig. 1A). Of note, a very small proportion (0.065%) of G4s may be associated with more than one cCRE. Therefore, in this study, we excluded these G4s when comparing the differences between various categories of G4s, such as the disparities in G4Hunter scores between PLS-G4s and pELS-G4s.

**Figure 1.**
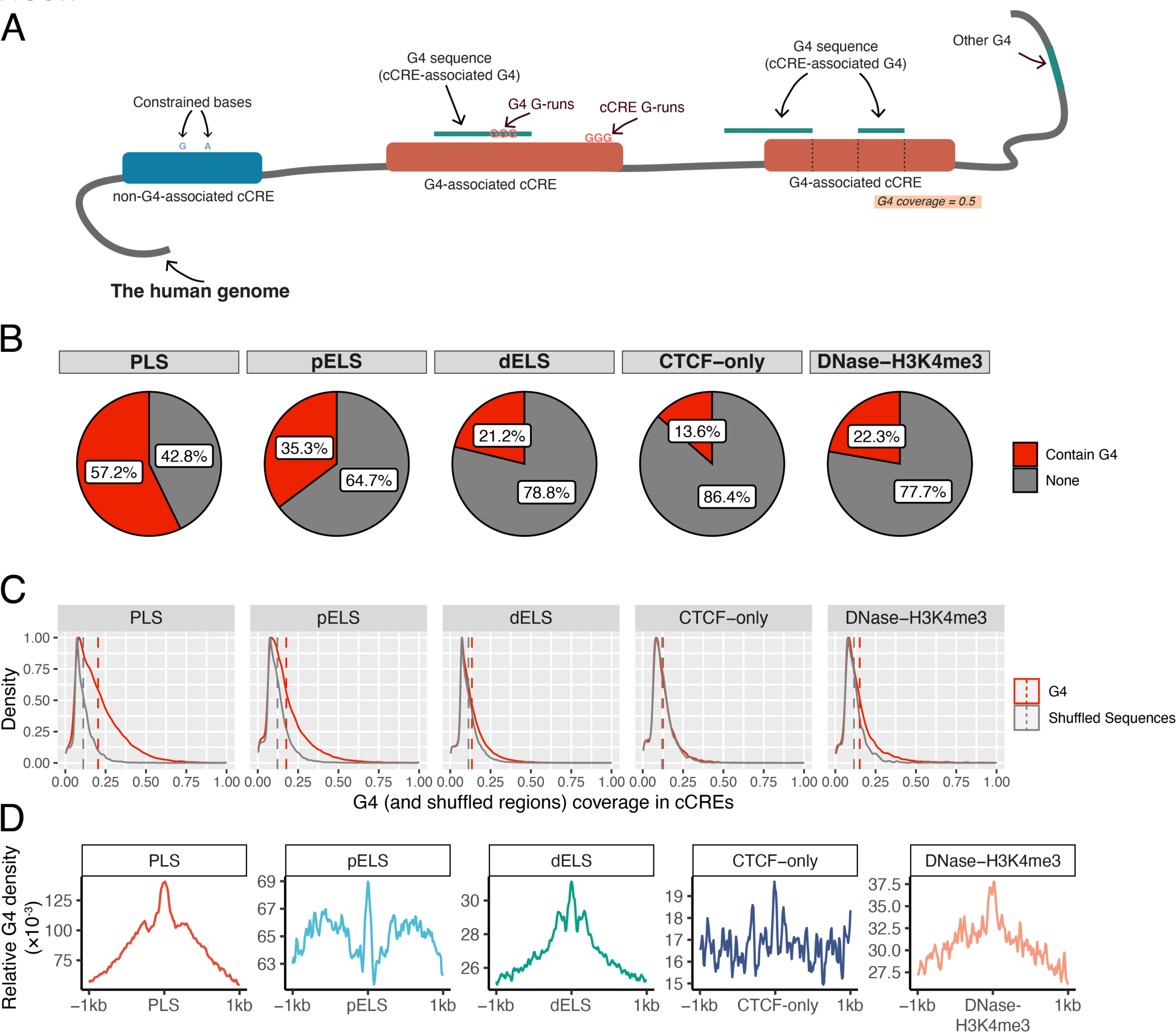
Overview of the enrichment patterns of G4s on cCREs. (**A**) Illustration of the definitions of G4-associated cCRE, cCRE-associated G4, *etc*. We define a cCRE that has an overlap with G4 as a G4-associated cCRE, and similarly, a G4 that has an overlap with a cCRE is defined as a cCRE-associated G4. The proportion of bases on a cCRE that participate in the composition of the G4 sequence is referred to as the coverage of G4 on that cCRE. (**B**) Pie charts illustrate the proportion of cCREs (PLS, pELS, dELS, CTCF-only, and DNase-H3K4me3 elements) that contain G4s or not. Red represents the proportion of cCREs that contain G4s, while gray represents the opposite. (**C**) Density plots show the differences of G4 coverage versus shuffled sequence coverage on different groups of cCREs. The coverage of G4 or shuffle sequences in each cCRE refers to the proportion of the cCRE sequence covered by G4 or shuffle sequences. Note that those cCREs that do not contain G4 or shuffled sequences were removed. Red and gray dashed lines indicate the average G4 coverage as well as the average shuffled sequence coverage on cCREs, respectively. (**D**) Distribution of G4s around cCREs. The x-axis indicates the relative genomic location (1kb upstream and downstream of the cCRE center), while the y-axis represents the relative density of G4s. Higher relative density signifies a greater enrichment of G4s in that region.

In individual analyses, we also used cell type-specific cCREs data, mainly for the K562 and the HepG2 cell lines, which were also obtained from the SCREEN database.

### Shuffled genomic regions

We utilized the shuffle function derived from the bedtoolsr package (V2.30.0-5) in R to randomly generate regions within the human genome with the same number and equal lengths as G4 sequences for comparative analysis, aiming to assess the enrichment significance of G4s in cCREs. In addition, we imposed restrictions on shuffle regions, excluding those that overlapped with the blacklist regions (https://github.com/Boyle-Lab/Blacklist/blob/master/lists/hg38-blacklist.v2.bed.gz) during the randomization process.

### Relative density calculation

In this study, we extensively calculated the relative density of B objects (*e.g.*, G4s) in both the upstream and downstream regions of A objects (*e.g.*, PLS elements). For instance, we examined the distribution of G4 sequences around different groups of cCREs, as well as the distribution of loop anchors upstream and downstream of G4-associated PLS elements and non-G4-associated PLS elements. All these calculations were performed using the EnrichedHeatmap package (V1.28.1) in R. We utilized the coverage mode of EnrichedHeatmap (V1.28.1) for calculation in all scenarios. The lengths of the upstream and downstream extensions, as well as the resolution of the density calculations, were adapted to the specific scenarios we analyzed, as outlined in the results section.

The principle of the calculation is as follows:

The regions upstream and downstream of the B objects are divided into non-overlapping windows with a width (resolution) of *L*. For the *i*-th window, the average G4 density *d_i_* was

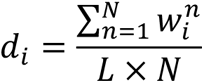

Where *N* represents the number of B objects, while *w* signifies the width that intersects with this window.

### TAD dataset and boundary definition

We extracted topologically associating domain (TAD) coordinates from the 3D Genome Browser database, using the hg38 version, for a variety of human cell lines and tissues. We defined TAD boundaries following the methods of Evonne *et al.* [19] we used in our previous study [15], which involved extending a specific width upstream from the TAD start coordinate and downstream from the TAD end coordinate. For instance, if the coordinates of a TAD were (chr1, X, Y), we then generated two boundary regions, with the coordinates of boundary 1 being (chr1, X-N, X) and the coordinates of boundary 2 being (chr1, Y, Y+N), where N is the length (width) of the extension. In this study, we examined the robustness of the results by testing TAD boundaries with widths of 5kb, 10kb, 20kb, and 50kb, respectively.

### Comparative analysis of cCRE enrichment at TAD boundaries

In this study, we compared the enrichment differences between CTCF-hybrid cCREs and CTCF-only cCREs at TAD boundaries. Assuming we have a set of TAD boundaries specific to a particular cell line or tissue, let the number of CTCF-only cCREs be represented as *k*. We first determined the count of CTCF-only cCREs that are located inside these boundaries and denoted this count as *m*. Considering that the number of CTCF-only cCREs is substantially smaller than that of CTCF-hybrid cCREs, we performed 1,000 random samplings. In each of these sampling, *k* samples were randomly selected from the CTCF-hybrid cCREs collection to form a subset. The number of elements in this subset located at TAD boundaries was signified as *n*. This process allowed us to estimate the background distribution of the number of CTCF-hybrid cCREs enriched at TAD boundaries when the quantity of CTCF-hybrid cCREs equaled that of CTCF-only cCREs, based on the results obtained from 1,000 random samplings.

We applied z-score to evaluate whether CTCF-hybrid cCREs are more enriched or depleted at TAD boundaries compared to CTCF-only cCREs, that is,

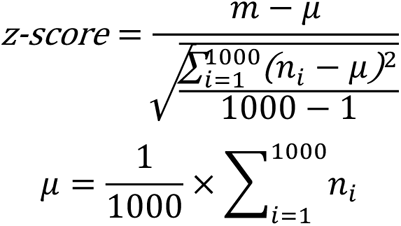

Where *m* is the total number of CTCF-only cCREs located inside TAD boundaries, whereas *n_i_* represents the total number of CTCF-hybrid cCREs located inside TAD boundaries in *i*-th sampling.

When the z-score is less than 0, it means that the enrichment level of CTCF-only cCREs at TAD boundaries is lower than the average level of CTCF-hybrid cCREs. Conversely, when it is greater than 0, it is higher.

### Cohesion-mediated chromatin loop dataset

The human pan-cell cohesion-mediated chromatin loop data was acquired from Fabian *et al.* [20] The genomic coordinates of loop anchors were then mapped to the hg38 assembly version using the liftOver software.

In accordance with different task scenarios, we computed and compared the relative density of loop anchors around CTCF-hybrid and CTCF-only cCREs. Additionally, we also examined the relative density of loop anchors around G4-associated cCREs and non-G4-associated cCREs. The calculation method for relative density have been described above.

### Comparison of epigenetic z-score values for different cCREs

The z-score matrix files for DNase, H3K4me3, H3K27ac, and CTCF epigenetic marks were gathered from the SCREEN database, which contains the raw signal data for each ENCODE sample used to identify cCREs. For each epigenetic mark (*e.g.,* DNase) and each cCRE group (*e.g.,* PLS), we examined the z-score differences of this epigenetic mark between G4-associated cCREs and other cCREs (non-G4-associated) in each sample separately. Subsequently, we computed the proportion of ENCODE samples in which the z-score values for the epigenetic marks of G4-associated cCREs were significantly higher than those of the other cCREs. We also determined the proportion of samples in which these values were lower or showed no statistically significant distinction.

### Transcription factor binding sites and motif data

We downloaded non-redundant human transcription factor binding site data from the ReMap2022 database (https://remap.univ-amu.fr). This dataset consists of a total of 68.6 M binding sites, with an average length of 313 bp. These binding sites were identified using various experimental techniques, including ChIP-seq, ChIP-exo, and others.

In addition, we collected human transcription factor motif data from Gregory *et al.* [21], resulting in an extensive collection of approximately 25.8 M motif instances. On average, these motif instances are around 11 bp in length. We refrained from merging motif instances derived from different transcription factors since this would obscure the binding preferences of multiple transcription factors for the same region.

We employed the EnrichedHeatmap package (V1.28.1) to compute the disparities in transcription factor binding density between G4-associated cCREs and other cCREs.

### DNA unmethylation data acquisition and intensity calculation

We obtained genome-wide unmethylated regions (blocks) of normal human tissues from Netanel *et al.* [22], where each of these blocks comprises a minimum of four CpGs, with the majority of CpGs within the block being identified as unmethylated. We used the liftOver software to convert the coordinates of unmethylated genomic blocks from the hg19 version to the hg38 version. We also obtained all of the blocks used in that study (regardless of whether they were methylated or not), which we filtered, retaining only the blocks containing at least four CpGs and employed them as background for comparisons.

First, for each type of tissue, we calculated the distribution density *D* of background blocks surrounding G4-associated cCREs and non-G4-associated cCREs (window width: 10 bp). Using the same approach, we subsequently computed the distribution density *d* of unmethylated blocks around them. Then the unmethylated intensities *I* of the window *n* around the two categories of cCREs in human tissue *t* can be computed as follows,

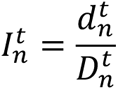

Where the determination of 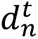 and 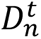 refer to the methods of relative density calculation above.

### Identification of G4 and cCRE G-runs

We implemented a customized script to identify G-runs within G4 sequences as well as within non-G4 sequences in cCREs. Typically, G4s with three or more quartets tend to be formed from sequences having at least the same number of consecutive guanines on each G-run, although there are exceptions (*e.g.,* sequences with bulges). Three-quartet G4s tend to be more stable than 2-quartets G4s; therefore, in this study, we only consider G-runs with at least 3 consecutive guanines. While some stable G-quadruplexes may be formed with interrupted G-runs (bulges), incorporating these possibilities into our computational analysis would introduce a significant level of complexity. Therefore, we do not take them into account here, aiming to simplify our analytical model.

We used regular expressions to detect G-runs in G4 sequences (G4s that overlapped with cCREs) on the forward strand. All potential G-runs in the forward strand cCREs were also identified, excluding those that intersected with forward strand G4s. Using the same approach, we also identified G4 G-runs and cCRE G-runs on the reverse strand of the human genome. Consequently, we can assess the differences between G4 G-runs and cCRE G-runs in terms of RegulomeDB regulatory scores, LINSIGHT scores, and other relevant factors.

### Acquisition and comparison of RegulomeDB, LINSIGHT, and gnomAD constraint scores

We devised four sets of comparative experiments to assess differences in the regulatory potential of regions associated with cCREs, as well as differences in the negative selection pressures to which they are subjected. These four sets include: (1) Comparing the differences in these metrics across different groups of G4s (*e.g.,* PLS-G4s, pELS-G4s, *etc.*). (2) Comparing the differences in these metrics across different group of cCREs (*e.g.,* G4-associated PLS versus non-G4-associated PLS, *etc.*). (3) Evaluating the differences in these metrics between G4 G-runs and the remaining sequences in G4s. Please note that the remaining sequences in G4s were obtained by subtracting G4 G-runs sequences from G4 sequences using the ‘subtract’ function provide by the bedtoolsr package (V2.30.0-5). (4) Examining the differences in these metrics between G4 G-runs and cCRE G-runs (excluding G4 G-runs).

We obtained the pre-calculated regulatory scores for common SNPs (dbSNP v153) with MAF ≥ 0.01 from the RegulomeDB website (https://regulomedb.org/regulome-search), which contains both probability scores and ranking scores, with the probability scores ranging from 0 to 1, where 1 represents the highest likelihood of being a regulatory variant, while the ranking scores taking the values from 1a to 7, where 1a indicates the one with the most supportive information and, therefore, potentially possess the strongest functional impact. We assigned these variants to different genomic regions for comparative analysis. For example, we evaluated the differences in regulatory probability scores as well as ranking scores among all potential variants on G4 G-runs and cCREs G-runs. If a certain region possesses a more pivotal role in regulatory functions, we can then observe that the regulatory scores in that region are generally higher.

We acquired the pre-calculated LINSIGHT score file (http://compgen.cshl.edu/LINSIGHT/LINSIGHT.bw) and mapped it to the hg38 assembly version (bigwig format). LINSIGHT scores can be used to estimate the extent of negative selection in non-coding regions of the human genome, such as the extent of negative selection differences between G4 G-runs and cCRE G-runs, with higher scores implying stronger negative selection. The bigWigAverageOverBed (V2) software was utilized to calculate the average negative selection strength for genomic regions such as G4 G-runs.

In addition, we acquired the human mutational constraint map from the gnomAD database (V3), which was imputed through the analysis of variations from ∼76k human genomes. However, due to the resolution of this constraint map being 1kb, we were unable to perform precise analysis on short sequences. Consequently, when evaluating disparities in gnomAD constraint scores (gnomAD uses z-scores to represent constraints. For simplicity, we will refer to them as ‘scores’ below), we only considered the distinctions between G4s located in different groups of cCREs, as well as the differences between G4-associated cCREs and non-G4-associated cCREs. Assuming that a genomic element (*e.g.,* a G4-associated PLS) overlap with *m* windows of this constraint map, and the widths (base pairs covered by this element) of the overlaps with these windows are (*w_1_*, *w*_2_,…, *w*_m_), the constraint scores for these *m* windows are (*s*_1_, *s*_2_,…, *s*_k_), then the constraint score for this element was calculated as,

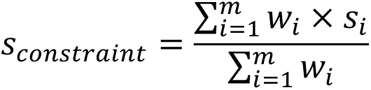

### phyloP conservation scores for 241 mammals and the definition of conversed bases

The phyloP conservation scores were obtained from the Zoonomia project (https://zoonomiaproject.org), which includes evolution-based conservation scores calculated for each position in the human genome. These scores were derived through the alignment of sequences from a comprehensive set of 241 mammalian species. We conducted comparisons of the differences in phyloP scores between different groups of G4s and between G4 G-runs and cCRE G-runs, *etc*. This analysis was carried out using the same methodology as for the assessment of LINSIGHT scores, as described above.

We also adopted the definition presented by Patrick *et al.* [23], defining bases with phyloP scores ≥ 2.27 as constrained bases. Using this threshold, we identified a total of 100,651,377 bases across the entire genome that are conserved during evolution.

### Distribution of constraint bases around G4s, cCREs, and G-runs

We compared the distribution of constrained bases around a certain group of cCREs (*e.g.,* PLS) and their associated G4s (*e.g.,* PLS-G4s). The calculations were performed using the EnrichedHeatmap package (V1.28.1), as detailed in the methods section above, with a resolution set to 10 bp.

Furthermore, we depicted the distribution of constrained bases in the upstream and downstream regions of G4 G-runs and cCRE G-runs located in distinct groups of cCREs regions. Given that the width of G-runs is usually very short, the resolution of the calculations here was increased to 1 bp. To estimate the distribution of constrained bases around G-runs, we conducted 100 random sampling experiments. In each random sampling, we independently selected 1,000 samples from both the G4 G-runs collection (*e.g.,* PLS G4 G-runs) and the cCRE G-runs collection (*e.g.,* PLS cCRE G-runs). We then calculated the relative density of constrained bases around the sampled samples. Finally, by analyzing the results from 100 random sampling experiments, we estimated the standard deviation errors in the distribution density of constrained bases around G4 G-runs and cCRE G-runs.

### Calculation of the proportion of G4s in activated cCREs

We calculated the proportion of endogenous G4s (as supported by G4 ChIP-seq experiments) and all G4s (without considering G4 ChIP-seq experiments) located on various types of activated cCRE elements for K562 and HepG2 cell lines, respectively. Activated cCREs are those that are supported by corresponding epigenetic signals (including DNase, H3K4me3, *etc*.) in specific cells. In contrast, quiescent cCREs are those lacking corresponding epigenetic data support in specific cells. For example, if a cCRE in the K562 cell line is found to be located in a heterochromatic region, then that cCRE is considered a quiescent cCRE.

As an example, the procedures for calculating the proportion of endogenous G4 located in activated PLS elements are as follows.

Due to the significant disparity in the quantities of the two types of G4s, we employed a sampling method to estimate the distribution of the proportions. The number of random samplings was set to 1,000. For each sampling, we randomly select 1,000 G4s from the set of endogenous G4s and calculated the number of these selected G4s that located on the active PLS elements; the number of selected G4s that located on quiescent PLS elements was also determined. The proportion of endogenous G4s located on activated PLS elements was then calculated as

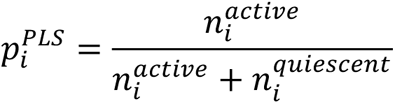

Where 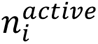 represents the number of endogenous G4s located in the activated PLS elements in the *i*-th sampling, while 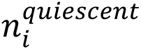 refers to the number of endogenous G4s located in the quiescent PLS elements.

The proportions of endogenous G4s on other types of activated cCREs, and the proportions of all G4s on activated cCRE elements were calculated with reference to the methods described above.

### Cell type-specific analysis

In this study, we compared the differences in transcription factor occupancy, DNA methylation, and chromatin accessibility between G4s supported by G4 ChIP-seq experiments and other G4s in the K562 and HepG2 cell lines.

The cell type-specific transcription factor binding data were sourced from the UCSC Genome Browser database (https://hgdownload.soe.ucsc.edu/goldenPath/hg38/encRegTfbsClustered/). We filtered all transcription factor binding sites for the K562 as well as the HepG2 cell line from this file, and in addition, the definition of transcription factors was strictly referenced to Matthew T. Weirauch *et al.* [24]. G4s predicted by G4Hunter software were divided into two categories based on whether they overlapped with G4 ChIP-seq experiments, and the coverage of various transcription factors on these two categories of G4s was calculated using bedtoolsr package (V2.30.0-5).

DNA methylation data and ATAC-seq data for the K562 and HepG2 cell lines were publicly obtained from the GEO database under accession number GSM2308596, GSM3633977, GSE170378 and GSE170251, respectively. We utilized the bigWigAverageOverBed (V2) software to calculate the average methylation and ATAC-seq signal values for the 200bp regions centered G4s.

### Pan-cancer data for chromatin accessibility, enhancer, and promoter activity

To investigate the activation status of G4-associated cCREs in cancer tissues, we acquired data on pan-cancer chromatin accessibility, enhancer, and promoter activity from publicly available databases, which were assessed using ATAC-seq, H3K27ac, and RNA-seq techniques. In this section of the study, we specifically focused on the cancer types for which all three types of data were available simultaneously.

The chromatin accessibility data were derived from the NIH Genomic Data Commons portal (https://gdc.cancer.gov/about-data/publications/ATACseq-AWG). In this study, cancer type-specific ATAC-seq peak datasets were utilized. We examined the coverage of ATAC-seq peaks for G4-associated cCREs and other cCREs, as well as the proportion of G4-associated cCREs and other cCREs located in open chromatin regions. Please note that chromatin accessibility is a necessary requirement for our analysis of pan-cancer enhancer data and promoter activity data. In other words, only the pan-cancer enhancers that overlap with ATAC-seq peaks will be included in subsequent analysis. The same requirement applies to both ELS (including pELS and dELS) elements and PLS elements. This definition is similar to the ENCODE project consortium definition of *cis*-regulatory elements [18].

We retrieved enhancer data identified through H3K27ac ChIP-seq in cancer tissues or cancer cell lines from the CenhANCER database [25] (https://cenhancer.chenzxlab.cn) with the typical enhancers were being considered for use. We assessed the distribution density of typical enhancers around enhancer-like cCREs (ELS), which encompass pELS and dELS.

The pan-cancer promoter activity was downloaded from the ICGC data portal. We utilized the raw promoter activity data from the PCAWG project, initially filtering out promoter activity data from normal samples within that dataset, retaining only cancer samples relevant to the cancer type under in our analysis. Subsequently, the PLS elements were assigned to the nearest promoters (TSS ± 200bp). The activity of a specific promoter for a given cancer type was defined as the average activity of that promoter across all samples of that cancer type. If a promoter overlapped with both G4-associated PLS elements and non-G4-associated PLS elements, it was excluded (∼12%) to ensure the accuracy of subsequent comparative analysis. We compared the differences in promoter activity between G4-associated PLS elements and non G4-associated PLS elements. Besides, we also evaluated the proportion of highly activated promoters associated with these two PLS element types, considering promoters with an activity exceeding 1.5 as highly activated [26].

## Results

### cCREs (candidate *cis*-regulatory elements) are enriched with G4s

To comprehensively characterize the relationship between G4s and *cis*-regulatory elements, we initially employed the G4Hunter software [8] to predict all potential G4 sequences in the human genome (see methods), and these predicted G4s will be used for subsequent analysis. In parallel, we retrieved the human candidate *cis*-regulatory elements (cCREs) from the SCREEN (Search Candidate *cis*-Regulatory Elements by ENCODE) database, in which human cCREs are primarily categorized into five groups, namely elements (or cCRE) with promoter-like signatures (PLS), proximal enhancer-like signatures (pELS), distal enhancer-like signatures (dELS), high DNase and CTCF signals only (CTCF-only), and high DNase and H3K4me3 signals only (DNase-H3K4me3). These cCREs are all defined based on epigenetic signals, including DNase-seq, H3K4me3, H3K27ac, as well as CTCF-only, and their distance from transcription start sites (see methods).

We first examined the proportion of cCREs overlapping with G4 motifs and compared it to the proportion overlapping with shuffled regions (see methods). As a result, more than half of the PLS elements (57.2%, Fig. 1B) overlap with G4 sequences, followed by pELS elements (35.3%, Fig. 1B). The dELS and DNase-H3K4me3 groups had around 20% of their elements overlapping with G4 sequences (Fig. 1B). The lowest overlap was observed in the CTCF-only group, with only 13.6% of its elements overlapping with G4 sequences (Fig. 1B). When compared to shuffled regions, we found that, except for the CTCF-only group, the overlap proportions of four out of five cCRE groups with G4 sequences were significantly greater than those with shuffled regions (Supplementary Fig. S1C), the exception being the CTCF-only elements for which G4 density is nearly identical to that of shuffled regions (13.6% and 13.9%, respectively).

We assessed G4 sequence and shuffled region coverage in cCREs separately, representing the proportion of bases covered by G4 or shuffled regions in each cCRE (Fig. 1A). Overall, across all five groups, there is a substantial number of cCREs with zero coverage of G4s or shuffled regions (Supplementary Fig. S2A). We observed that there is a significant difference in the average coverage of G4 sequences within the PLS group compared to shuffled regions (Supplementary Fig. S2A), while in the CTCF-only group, almost no difference could be found (Supplementary Fig. S2A). This discrepancy exists when we removed the cCREs that were not covered by G4s or shuffled regions. Without considering the CTCF-only group, we still found significant differences in coverage between G4s and shuffled regions (Fig. 1C, Supplementary Fig. S2B), particularly in the PLS and pELS groups (Fig. 1C, Supplementary Fig. S2B), suggesting that the higher coverage of G4s in cCREs cannot be simply explained by the fact that more cCREs were overlapped with G4s. To further valid this, we analyzed the distribution pattern of G4s around cCREs. A significant enrichment of G4 sequences in the vicinity of the cCRE center (less than 100 bp) was observed (Fig. 1D), whereas, in contrast, no specific distribution pattern was found in shuffled regions surrounding cCREs (Supplementary Fig. S3). Besides this, we found that the G4s in PLS and pELS groups tended to have higher absolute G4Hunter scores (Supplementary Fig. S4), indicating that G4s in these elements possessed greater stability.

### G4s are enriched at CTCF-bound cCREs accompanied by H3K4me3 or H3K27ac signals

We and others previously reported that G4s are tightly associated with the binding of CTCF, with CTCF binding sites significantly enriched around G4s [27–29]. However, in this study, our analysis showed a weak correlation between G4s and CTCF-only cCREs, which seems to challenge the existing conclusions and findings. Indeed, other groups of cCREs may also contain CTCF binding (CTCF-bound) signals that, together with enhancer (H3K27ac) or promoter (H3K4me3) signals, form the basis of PLS and ELS elements. To elucidate the precise association patterns between G4s and CTCF binding elements, we proceeded to perform comparative analysis between CTCF-only cCREs, which exclusively contain CTCF-bound signals, and cCREs containing both CTCF and H3K4me3 or H3K27ac signals (referred to as CTCF-hybrid cCREs or CTCF-hybrid elements for simplicity, Fig. 2A).

**Figure 2.**
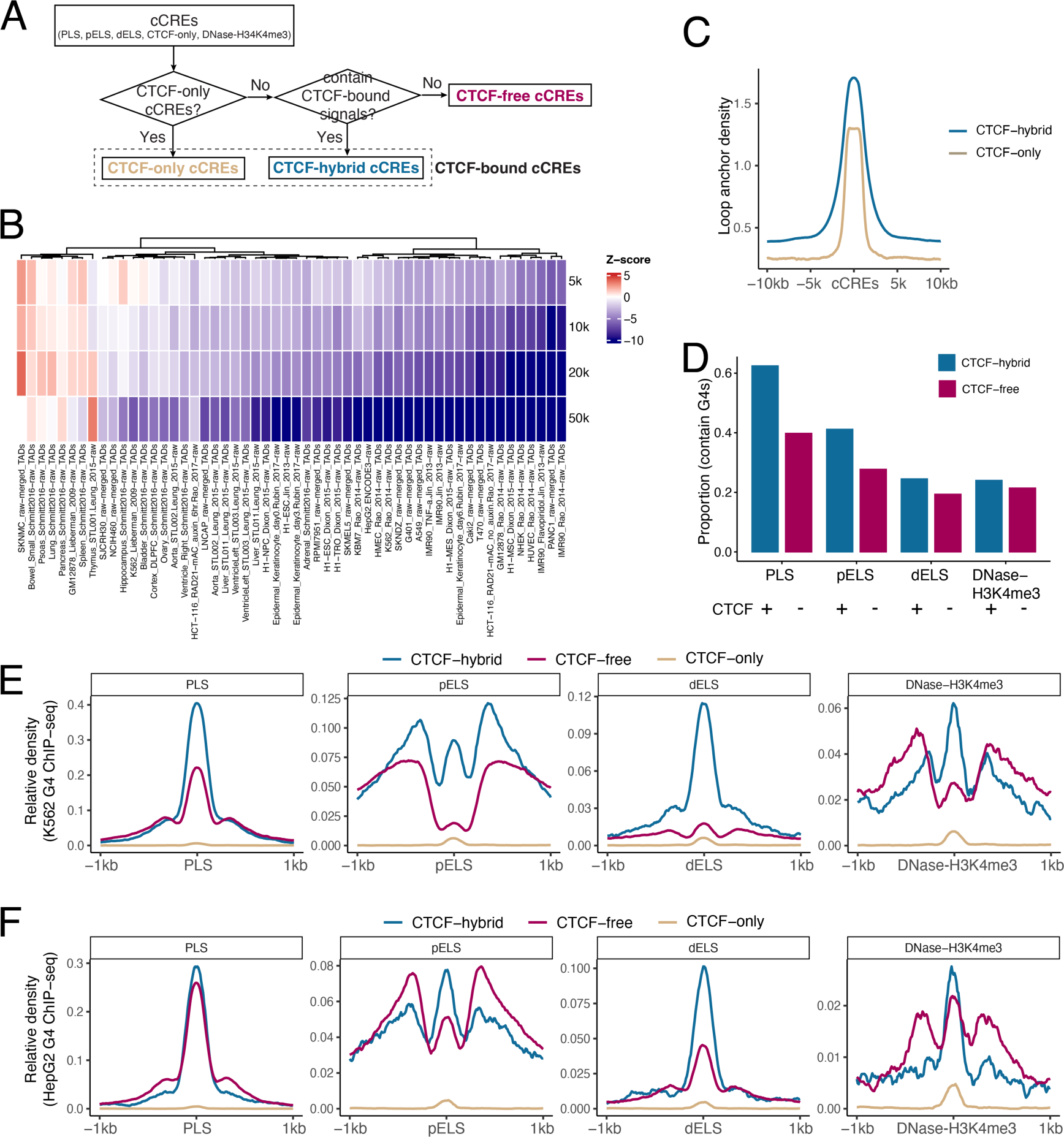
Association mode of G4s with cCREs exhibiting different CTCF occupancy patterns. (**A**) Flowchart illustrates our approach to categorize cCREs based on the occupancy patterns of CTCFs. (**B**) Z-score values for the enrichment of CTCF-only cCREs compared to CTCF-hybrid cCREs on TAD boundaries of different sizes (5, 10, 20 or 50 kb) in different cells or tissues. Red cell indicates that the enrichment level of CTCF-only cCREs at that TAD boundary is higher than CTCF-hybrid cCREs, while blue cell indicates a lower enrichment level. (**C**) Relative density of chromatin loops aound CTCF-hybrid and CTCF-only cCREs, with blue and brown curves signify CTCF-hybrid and CTCF-only cCREs, respectively. (**D**) Proportion of CTCF-hybrid and CTCF-free cCREs containing G4s. (**E-F**) Relative density of G4 ChIP-seq high-confidence peaks around CTCF-hybrid and CTCF-free cCREs for K562 (**E**) and HepG2 (**F**) cell lines. The relative density G4 ChIP-seq high-confidence peaks around CTCF-only cCREs was also put on the same scale for comparison.

Considering that G4s are preferentially located at the boundaries of topologically associating domains (TADs) [27], we therefore suspected that the distribution of CTCF-only and CTCF-hybrid elements at TAD boundaries would differ. We compared the enrichment differences of CTCF-only and CTCF-hybrid elements at TAD boundaries across various cell lines or tissues (see methods). As expected, the enrichment level of CTCF-hybrid elements was higher than that of CTCF-only elements in most of the cell lines or tissues (Fig. 2B). A similar phenomenon was found when we focused on cohesin-mediated anchor regions of chromatin loops (Fig. 2C). That is, CTCF-hybrid elements are more enriched at the anchor regions of chromatin loops compared to CTCF-only elements (Fig. 2C). We speculate that the differential enrichment distribution of CTCF-hybrid elements in specialized regions of higher-order chromatin structure, may facilitate a closer association with G4s, as compared to CTCF-only elements.

In addition, we separately calculated the distribution of G4 sequences around CTCF-hybrid and CTCF-only cCREs. CTCF-hybrid cCREs are significantly more enriched with G4 sequences in their vicinity than CTCF-only cCREs (Supplementary Fig. S5A), indicating a clear association between G4 and CTCF binding. However, the simultaneous presence of enhancer or promoter signals in CTCF-hybrid cCREs made it challenging to rule out whether the association of G4s with CTCF-hybrid cCREs might be due to enhancers or promoters alone. Therefore, to further confirm the independence of the relationship, we compared the differences in the proportions of CTCF-hybrid and CTCF-free cCREs overlapped with G4s. The results showed that a higher proportion of cCREs are associated with G4s when they simultaneously contain CTCF signals, which implies a genuine positive close connection between CTCF binding and G4 localization, but only for the PLS and pELS cCRE groups (Fig. 2D). Of note, the G4 density around CTCF-hybrid cCREs tended to be the highest (Supplementary Fig. S5B).

Since the association between CTCF binding and G4 localization can be largely explained by CTCF binding preference for G4 structures [28, 29], we subsequently conducted cell type-specific analysis to investigate whether there are any differences in the occupancy of G4 structures around cCREs with different CTCF binding states. We obtained cCREs for the K562 and HepG2 cell lines from the SCREEN database and, in addition, retrieved high-confidence G4 ChIP-seq peak data for these two cell lines (see methods). Similarly, we divided the elements in different cCRE groups into CTCF-hybrid and CTCF-free subgroups and calculated the distribution of G4 ChIP-seq peaks around these subgroups. We also calculated the distribution of G4 ChIP-seq peaks around CTCF-only cCREs for comparison. Indeed, we observed an enrichment of the G4 structures around the CTCF-only cCREs (Supplementary Fig. S5C-D), but this enrichment almost disappeared when we compared the CTCF-only cCREs with other cCREs at the same scale (Fig. 2E-F), indicating that the enrichment of the G4 structures around the CTCF-only cCREs is much less significant than that around the other types of cCREs. Moreover, the average distribution density of the G4 secondary structures around these cCREs tends to be higher when CTCF binding signals are present (Fig. 2E-F), suggesting that the co-occurrence of CTCF and H3K4me3 or H3K27ac signals is associated with a stronger G4 structure occupancy, and G4 structures do not strongly preferentially present in open chromatin regions that contain only CTCF signals.

### cCRE-G4s are inclined to possess regulatory significance and are under negative selection in the human genome

The preferential distribution of G4s on cCREs raises a question: could variants in cCRE-G4s potentially affect the regulatory functions of the genome? We then conducted a comprehensive investigation into the regulatory potential of all variants that can be mapped to G4s or cCREs regions, using RegulomeDB as a reference [30]. First, we found that the variants on PLS-G4s exhibited the highest RegulomeDB scores (Fig. 3A), followed by pELS-, dELS-, and CTCF-only-G4s, while the scores for variants on DNase-H3K4me3-G4s were relatively lower (Fig. 3A). Besides, the regulatory variation scores on cCRE-G4s are significantly higher than on other G4s, suggesting the potential significant regulatory roles of cCRE-G4s. This observation remains valid when examining the RegulomeDB ranking scores in these G4 regions, where cCRE-G4s showed a higher proportion of high-ranking scores (Supplementary Fig. S6A). Furthermore, we inspected whether the regulatory scores of variants in cCREs associated with G4s or not would show differences. We found that variants in G4-associated cCREs displayed higher regulatory scores and higher proportions of high-ranking scores (Supplementary Fig. S6B-C; Wilcoxon test, *P* = 1.76×10^-70^, *P* = 1.04×10^-210^, *P* = 0, *P* = 1.27×10^-86^, and *P* = 3.42×10^-62^ for comparisons in the PLS, pELS, dELS, CTCF-only, and DNase-H3K4me3 groups, respectively), which implies that the presence of G4s may enhance the regulatory potential of their corresponding cCREs.

**Figure 3.**
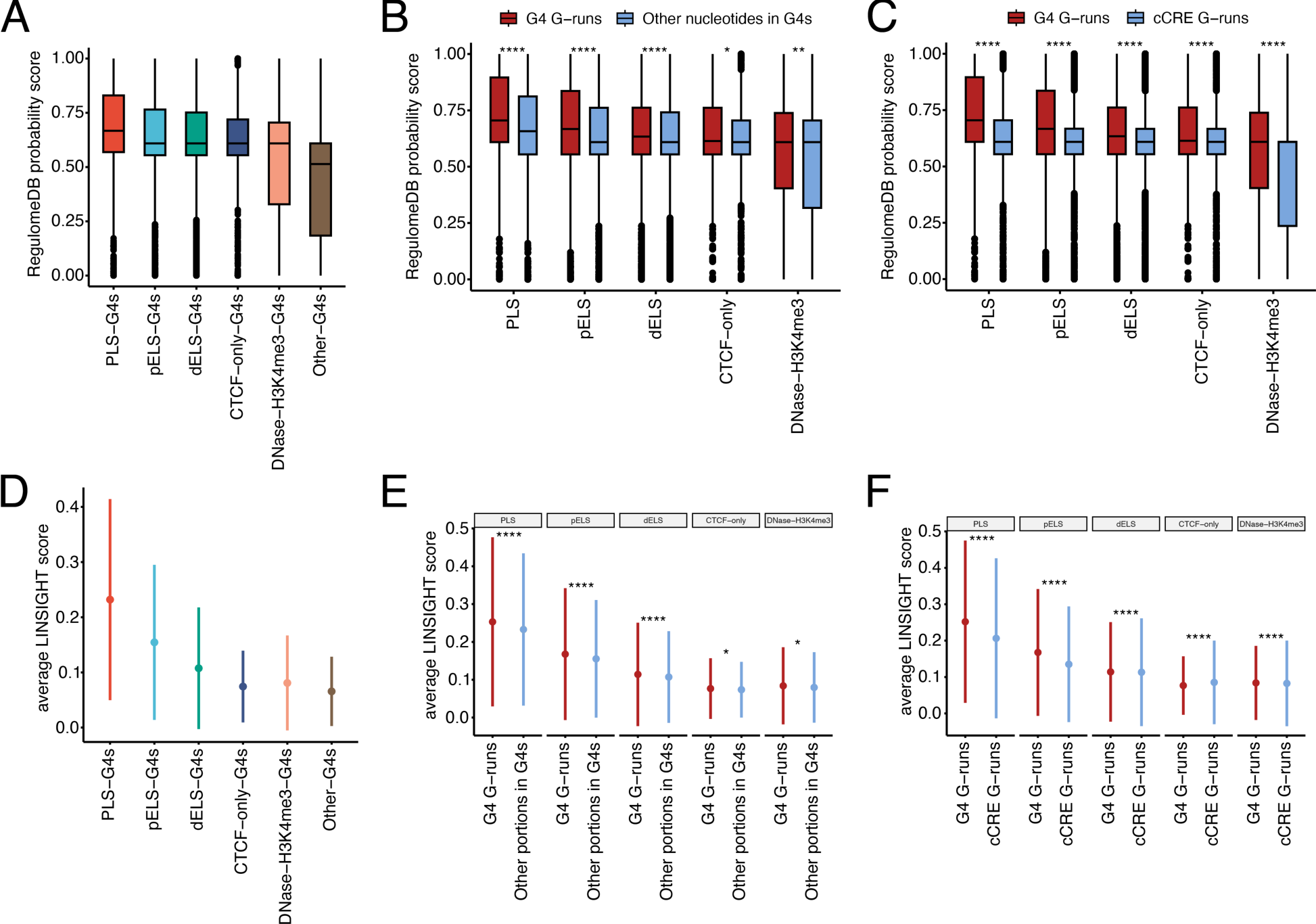
Overview of RegulomeDB and LINSIGHT scores for G4s and cCREs. (**A**) The RegulomeDB probability scores of variants located in different groups of G4s. The y-axis shows the RegulomeDB probability scores for all variants assigned to G4s located in different groups of cCREs. (**B**) Boxplot showing the differences in RegulomeDB probability scores for variants located in G4 G-runs versus variants in other nucleotides of G4s (the remaining nucleotides that are not contribute to G-runs) across five groups of cCREs. Wilcoxon test, *: *P* < 0.05, **: *P* < 0.01, ****: *P* < 0.0001. (**C**) Similar to (**B**), with the distinction that we consider the differences in RegulomeDB probability scores for variants on G-runs of G4s and cCREs. Wilcoxon test, ****: *P* < 0.0001. (**D**) The average LINSIGHT scores for different groups of G4s. The mean and the standard deviation error limits are visualized by error plots. (**E**) The average LINSIGHT scores for G4 G-runs and the remaining parts of G4s excluding G-runs (other portion) in all five groups of cCREs. The red and blue error plots indicate the mean and the standard deviation error limits of the average LINSIGHT scores for G4 G-runs and the other portion in G4s. Wilcoxon test, *: *P* < 0.05, ****: *P* < 0.0001. (**F**) Resembling (**E**), with the comparison conducted between G4 G-runs and cCRE G-runs. Wilcoxon test, ****: *P* < 0.0001.

Given the important role of G-runs for the stability of the G4 sequence, we next compared the regulatory scores of variants harbored on G-runs within G4 sequences with those in the remaining sequences of G4s. Generally, nucleotides within runs of three or more G are more likely to be involved in the formation of a G-tetrad, although this is not absolute since the actual formation of G-quadruplex secondary structures may involve complex sequence patterns. We found that the variants on PLS and ELS G-runs present higher regulatory scores compared to the remaining variants in G4 sequences (Fig. 3B; Wilcoxon test, *P* = 8.07×10^-29^, *P* = 4.55×10^-32^, *P* = 1.58×10^-51^, *P* = 3.61×10^-2^, and *P* = 9.25×10^-3^ for comparisons in the PLS, pELS, dELS, CTCF-only, and DNase-H3K4me3 groups, respectively), while also having a higher proportion of 1a ranking scores (see methods; 1a corresponding to the category with the highest likelihood being functional; Supplementary Fig. S6D). However, the regulatory potential of variants on G4 sequences, especially those on G-runs, may be solely due to the compositional biases of nucleotide types. To eliminate this effect, we proceeded to analyze the regulatory score differences of variants on G-runs derived from G4s and those originating from cCREs that are not involved in the composition of G4 sequences (see methods). The results revealed that variants on the G-runs of G4s exhibited higher regulatory scores compared to those on cCRE G-runs (Fig. 3C; Wilcoxon test, *P* = 1.94×10^-130^, *P* = 1.82 ×10^-166^, *P* = 0, *P* = 7.27×10^-9^, and *P* = 1.78×10^-7^ for comparisons in the PLS, pELS, dELS, CTCF-only, and DNase-H3K4me3 groups, respectively), along with a higher proportion of 1a ranking scores (Supplementary Fig. S6E). This suggests that variants occurring on G-runs of G4 sequences have a greater potential impact on genome regulation, while the functionality of G4s cannot be simply explained by their high G/C content.

Since G4s are potential functional elements within human cCREs, are they less prone to mutation in the human genome? With this question, we then assessed whether cCRE-G4s in the human genome are under negative selection using the similar approach as mentioned above, but with the assistance of LINSIGHT scores (see methods) [31]. First, we observed that G4s located in the PLS and pELS regions displayed relatively higher average LINSIGHT scores (Fig. 3D), suggesting that G4s in these regions are under stronger negative selection, making variants in these G4 regions more likely to be deleterious. Intriguingly, we found that PLS and ELS (pELS and dELS) cCREs containing G4 sequences also had higher average LINSIGHT scores (Supplementary Fig. S7; Wilcoxon test, *P* = 0, *P* = 0, and *P* = 0 for comparisons in the PLS, pELS, and dELS groups, respectively), implying that the presence of G4 sequences may render these regions more susceptible to negative selection rather than neutral selection, which could be intertwined with the potential functional role of G4s, despite the fact that the average LINSIGHT scores for G4-associated cCREs in both CTCF-only and DNase-H3K4me3 groups were slightly lower compared to non-G4-assciated cCREs (Supplementary Fig. S7; Wilcoxon test, *P* = 1.10×10^-3^ and *P* = 3.12×10^-13^ for comparisons in the CTCF-only, and DNase-H3K4me3 groups, respectively). Besides, the average LINSIGHT scores for G4 G-runs were also greater than those of the remaining part of the G4 sequences (Fig. 3E; Wilcoxon test, *P* = 9.46×10^-58^, *P* = 1.64×10^-39^, *P* = 8.36×10^-18^, *P* = 3.76×10^-2^, and *P* = 2.47×10^-2^ for comparisons in the PLS, pELS, dELS, CTCF-only, and DNase-H3K4me3 groups, respectively). We propose that G4 G-runs play a critical role in stabilizing these secondary structures and are therefore subject to higher constraints in the human genome.

To further confirm that the stronger negative selection on G4 G-runs was not simply caused by high G/C content, we then examined the average LINSIGHT scores of G-runs in cCREs that are not involved in G4 sequence composition. Consequently, we observed that the average LINSIGHT scores of the G4 G-runs were significantly higher than those of the cCRE G-runs in the PLS, ELS, and DNase-H3K4me3 groups (Fig. 3F; Wilcoxon test, *P* = 0, *P* = 0, *P* = 0, *P* = 1.65×10^-231^ for comparisons in the PLS, pELS, dELS, and DNase-H3K4me3 groups, respectively), indicating that the heightened negative selection on G4 G-runs was not merely due to the nucleotide composition of G4 sequences, though we noted that the average LINSIGHT scores for G4 G-runs in the CTCF-only group were lower (Fig. 3F; Wilcoxon test, *P* = 2.63×10^-60^).

Finally, we also used the human genome-wide mutational constraint map from the gnomAD (Genome Aggregation Database) database [32], which aggregates large-scale human variants data, for further quantification. Due to the resolution limit of the constraint map (1kb), we were unable to accurately compute and compare the constraints on regions with shorter widths, such as G4 G-runs and cCRE G-runs. Therefore, we no longer discuss the gnomAD constraint differences between them. Overall, the G4s in PLS and pELS presented higher constraint z-score values (Supplementary Fig. S8A), which indicates that the observed number of variants in these G4 regions was lower than expected. This is consistent with the results based on the LINSIGHT model, except that the gnomAD constraint z-scores were derived from a substantial amount of human genomic mutation data. Moreover, we found that those cCREs containing G4 sequences showed higher constraint z-score values as well (Supplementary Fig. S8B; Wilcoxon test, *P* = 1.87×10^-230^, *P* = 0, *P* = 0, *P* = 1.14×10^-95^, and *P* = 2.47×10^-148^ for comparisons in the PLS, pELS, dELS, CTCF-only, and DNase-H3K4me3 groups, respectively).

In general, these findings suggest that the presence of G4s makes the cCRE sequences more constrained, potentially attributable to the significant genomic regulatory functions associated with G4s.

### G4 G-runs in cCREs near the transcription start site are highly constrained across 241 mammalian genomes

We obtained the phyloP scores from the Zoonomia Project database (https://zoonomiaproject.org/), which are derived from the alignment of 241 mammalian genomes. These scores were then used to estimate the evolutionary constraints on G4s, as well as cCREs, and to determine whether G4s in cCREs were under selective pressures during evolution. Similar to our aforementioned findings, G4s in PLS elements are under greater selection pressure, followed by the G4s in pELS. In contrast, the remaining three groups of G4s and those G4s that are not associated with cCREs undergo lower selection pressure (Fig. 4A). Moreover, the average phyloP scores of G4 G-runs tend to be higher than the remaining sequences in G4s (Fig. 4B; Wilcoxon test, *P* = 0, *P* = 0, *P* = 0, *P* = 1.13×10^-187^, and *P* = 0 for comparisons in the PLS, pELS, dELS, CTCF-only, and DNase-H3K4me3 groups, respectively). Nevertheless, when we focus on the comparison of average phyloP scores between G4s and cCREs, we found that although the average phyloP scores of G4s on pELS are higher than those on pELS cCREs, for the other three groups of cCREs (including dELS, CTCF-only, and DNase-H3K4me3 groups), the average phyloP scores of G4s on them are lower (Supplementary Fig. S9; Wilcoxon test, *P* = 3.14×10^-1^, *P* = 8.46×10^-87^, *P* = 0, *P* = 0, and *P* = 5.01×10^-268^ for comparisons in the PLS, pELS, dELS, CTCF-only, and DNase-H3K4me3 groups, respectively). A similar phenomenon was observed when we analyzed the average phyloP scores of the G4 G-runs and the cCREs G-runs that do not comprise G4 sequences (Fig. 4C). Specifically, the average phyloP score of G4 G-runs in PLS and pELS was higher than that of G-runs in cCREs (Fig. 4C; Wilcoxon test, *P* = 0 and *P* = 9.05×10^-60^ for comparisons in the PLS and pELS groups, respectively). However, in the other three groups of cCREs, the situation is reversed (Fig. 4C; Wilcoxon test, *P* = 0, *P* = 4.39×10^-106^, and *P* = 5.88×10^-66^ for comparisons in the dELS, CTCF-only, and DNase-H3K4me3 groups, respectively). Since PLS and pELS are elements located relatively close (<2kb) to the transcription start site (TSS), we suspect that those G4s further away from the TSS may not be as conserved compared to cCREs during the process of evolution.

**Figure 4.**
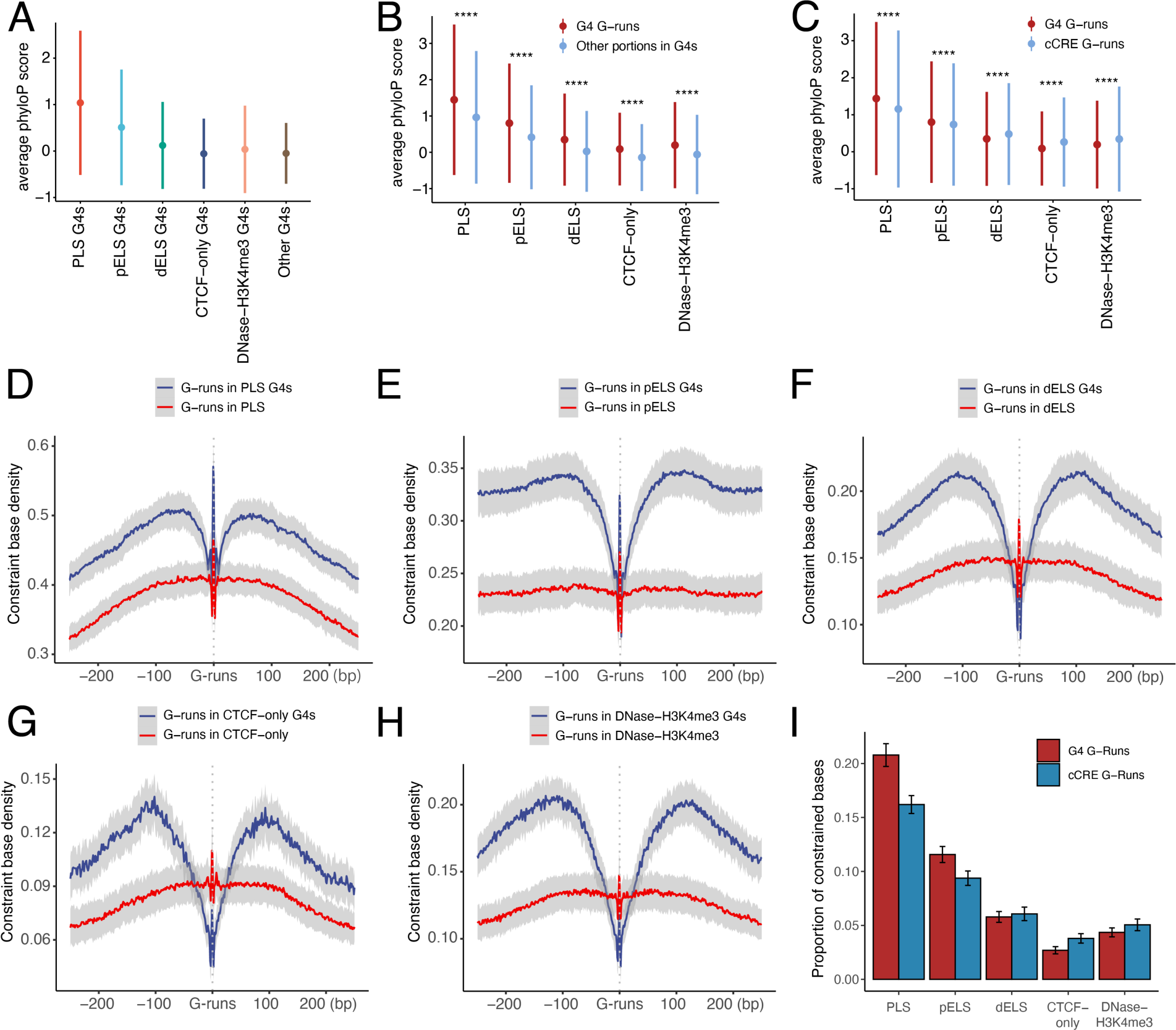
Assessing the extent of constraints on cCRE-G4s in mammalian evolution. (**A**) The average phyloP scores for different groups of G4s, with dots and ranges indicate the mean and standard deviation errors, respectively. (**B-C**) The differences in average phyloP scores between G4 G-runs and other portions in G4s across the five types of cCREs (**B**). While in (**C**), the error plots depict the distinctions between G4 G-runs and cCRE G-runs in different cCRE groups. Wilcoxon test, ****: *P* < 0.0001. (**D-H**) The distribution pattern of constraint bases around G4 G-runs and cCRE G-runs in PLS, pELS, dELS, CTCF-only, and DNase-H3K4me3 cCREs, respectively. Blue and red curves denote the constraint base density surrounding G4 G-runs and cCRE G-runs, respectively. (**I**) Proportion of G-runs in G4s and cCREs covered by constraint bases across all five groups of cCREs. The error bars correspond to the standard deviations.

To further confirm this, we first defined bases with phyloP scores ≥ 2.27 as constrained bases, using the threshold proposed by Patrick *et al*. [23]. Subsequently, we calculated the distribution of constrained bases upstream and downstream of G4s that overlapped with cCREs, as well as the distribution of constrained bases around cCREs. Surprisingly, the constrained bases exhibited a localized depletion around cCRE-G4s, while they displayed an enriched distribution in the vicinity of cCRE centers (Supplementary Fig. S10A-E). Considering that the bases within G4 sequences that are not part of G4 G-runs are likely to be less conserved, we proceeded to investigate the distribution of constrained bases centered on both G4 G-runs and cCRE G-runs, respectively. Our findings revealed that G-runs represent highly constrained regions, and this holds true for G4 G-runs as well (Fig. 4D-H). That is, the G-runs in G4s are more conserved compared to the remaining sequences of G4s (Fig. 4D-H). Apart from this, we also noticed that G4 G-runs are more constrained than cCREs G-runs in PLS and pELS elements (Fig. 4D-H), while the opposite is true in the remaining three groups of cCREs (Fig. 4D-H), which is consistent with our analysis above. The similar phenomenon was observed when we calculated the proportion of constrained bases in G4 G-runs and cCRE G-runs (Fig. 4I), which strongly suggests that the G-runs in G4s near the transcription start site are highly constrained in mammals.

### The presence of G4s is associated with a favorable local environment for *cis*-regulatory elements

Previous studies have shown that the presence of G4s is associated with epigenetic marks [33, 34]. We wondered whether the occupancy of G4s in cCREs is also linked to a local environment that is more favorable for the activation or functioning of cCREs. Since the functions of promoters and enhancers are relatively clear, in this section, we only consider PLS and ELS elements.

We first divided the cCREs into two subgroups based on whether they overlapped with G4s or not, and then compared the z-score signal values of DNase, H3K4me3, H3K27ac, and CTCF in these two subgroups across all samples utilized in the SCREEN database. We found that G4-associated cCREs exhibited higher DNase signal values in most samples, irrespective of the cCRE type (Fig. 5A). G4-associated PLS cCREs displayed significantly higher H3K4me3 and H3K27ac z-score signal values in nearly all samples compared to other PLS cCREs not associated with G4s (Fig. 5A). This is also the case in pELS, with the exception that, in a small subset of samples, the H3K27ac signals of G4-associated cCREs were lower than that of other cCREs (Fig. 5A). In most samples, both H3K4me3 and H3K27ac signal values of G4-associated cCREs were significantly greater than those of other cCREs, which was roughly similar to the proportions observed in CTCF signal values (Fig. 5A). Nevertheless, in most samples, we do observe higher values of these epigenetic marks in G4-associated cCREs compared to other cCREs.

**Figure 5.**
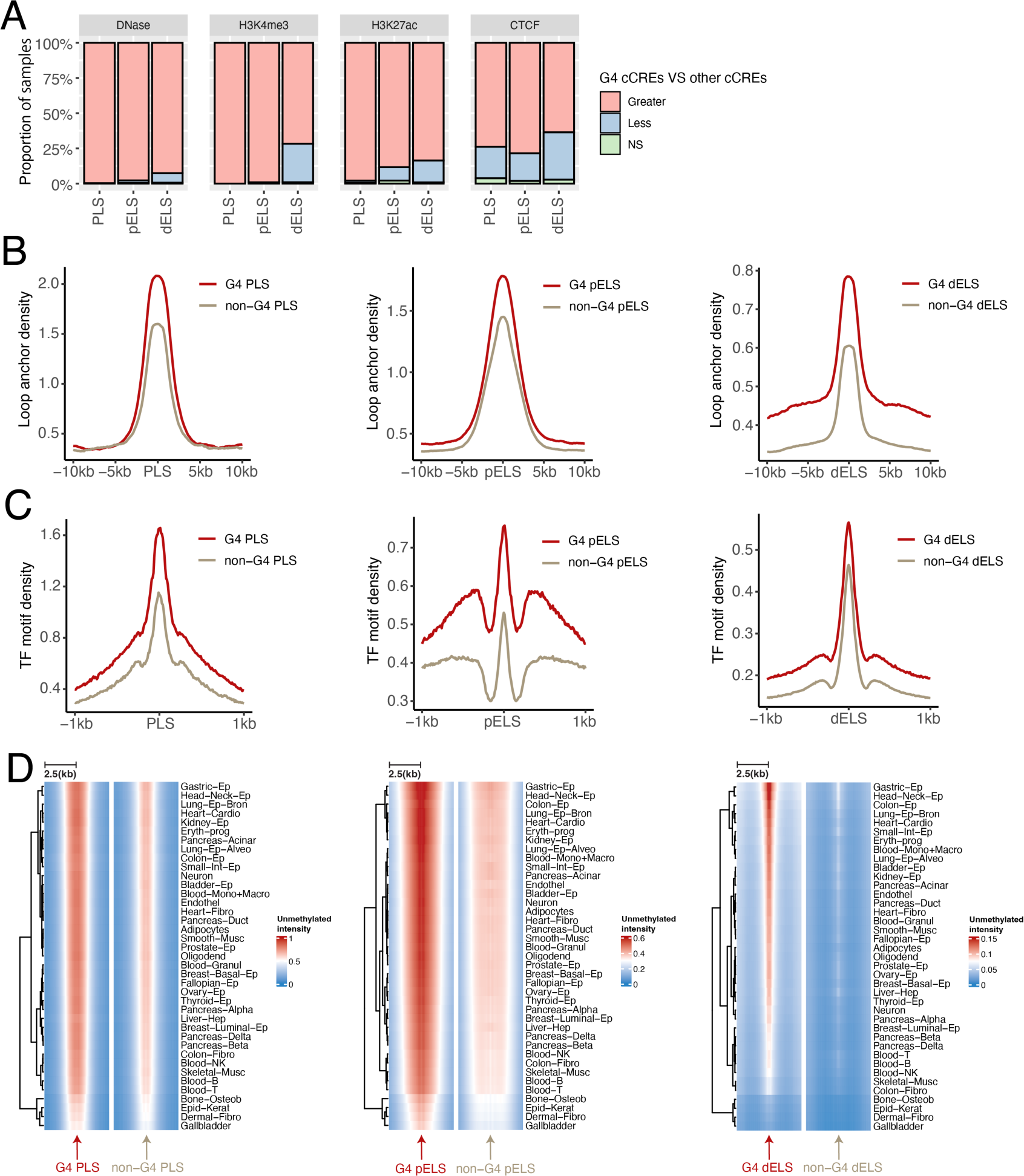
Distribution pattern of epigenetic marks surrounding the cCREs associated with G4s or not. (**A**) Proportion of the samples used in the SCREEN database that exhibit z-score differences in DNase, H3K4me3, H3K27ac, and CTCF signals between the two subtypes of cCREs associated with G4s or not. Pink bar represents the proportion of samples in which the signal value of G4-associated cCREs on that epigenetic mark is significantly greater than that of other cCREs (non-G4-associated cCREs); while blue bar indicates the opposite. Green represents the proportion of samples where there is no difference between the two subgroups. PLS, pELS and dELS cCREs are considered separately. (**B**) Loop anchor distribution around different groups of cCREs (PLS, pELS, dELS). The red curve represents the density of loop anchors around G4-associated cCREs, while the brown curve shows the density of loop anchors around non-G4-associated cCREs. (**C**) Similar to (**B**), except that the density of the TF binding motif is calculated. (**D**) Heatmap of the unmethylated intensity around G4-associated and non-G4-associated PLS, pELS, dELS cCREs in diverse tissues with red blocks represent higher unmethylated intensity, while blue blocks indicate lower unmethylated intensity.

Given that enhancers can interact with promoters through spatial proximity, thereby regulating gene expression, we were intrigued by the potential differences in the distribution of chromatin loop anchor regions around the two cCRE subgroups. For this purpose, we retrieved data on human pan-cell type cohesin-mediated chromatin loops [20] and calculated the distribution densities of loop anchors around the two subgroups of cCREs associated with G4s or not. As expected, G4-associated cCREs tended to be preferentially distributed in anchor regions compared to other cCREs (Fig. 5B). This is particularly pronounced in dELS cCREs, indicating that the presence of G4s may enhance the regulation of gene expression by cCREs through three-dimensional mechanisms, which is in line with our previous findings [27]. Additionally, using a similar approach, we also calculated the distribution density of transcription factor binding sites around the two subgroups of cCREs. We utilized the motif instances derived from the study of Gregory *et al.* [21] and the transcription factor binding sites sourced from the ReMap2022 database [35] for our calculations. The motif instances of transcription factors exhibited a relatively higher density around G4-associated cCREs (Fig. 5C), and the same trend was also observed when analyzing the transcription factor binding sites data provided by the ReMap2022 database (Supplementary Fig. S11A-C), suggesting that the presence of G4s in cCREs may enhance the interaction between transcription factors and cCREs, which coincides with a previous study indicating that G4s serve as hubs for transcription factor binding [36]. Considering that DNA methylation can influence the activity of cCREs [3], we next examined whether the occurrence of G4s might be potentially associated with a lower methylation landscape in cCREs. In this regard, we characterized the intensity of genome-wide unmethylated regions in human cells in the vicinity of cCREs in both subgroups. As a result, there was a stronger hypomethylation around G4-associated cCREs in most of the cell types (Fig. 5D), which was particularly pronounced in PLS as well as pELS elements, suggesting that the presence of G4s may lead to a low methylation state in cCRE regions, making them more susceptible to activation and utilization.

### The G4 structure itself, and not simply its associated G-rich sequence, is associated with cCRE activation

To confirm whether it is the G4 secondary structure or the corresponding G-rich sequence that is associated with the activation of cCREs, we then parsed the pattern of linkage between cell-type specific G4s and cCREs in the K562 and HepG2 cell lines. We initially evaluated the proportion of G4s supported by G4 ChIP-seq experiments (hereafter referred to as endogenous G4s) that are located on activated cCREs (*e.g.,* PLS, see methods). Besides, the background proportion was also estimated without considering whether G4s are supported by ChIP-seq experiments or not (see methods). As a result, we found that for endogenous G4s, they are overwhelmingly located on activated cCREs rather than quiescent cCREs, which is particularly evident for G4s on PLS elements both in the K562 and HepG2 cells (Fig. 6A). Interestingly, endogenous G4s are selectively enriched around activated cCREs, with no apparent enrichment observed around quiescent cCREs (Supplementary Fig. S12A-B). In contrast, those G4s that do not form secondary structures exhibit an association with cCREs independent of their activation status (Supplementary Fig. S12A-B). For instance, in K562 cells, we observed the enrichment of those G4s around both activated and quiescent PLS regions (Supplementary Fig. S12A). This suggests that the activation of cCREs cannot be determined by G4-compatible sequences alone and potentially requires the formation of secondary structures.

**Figure 6.**
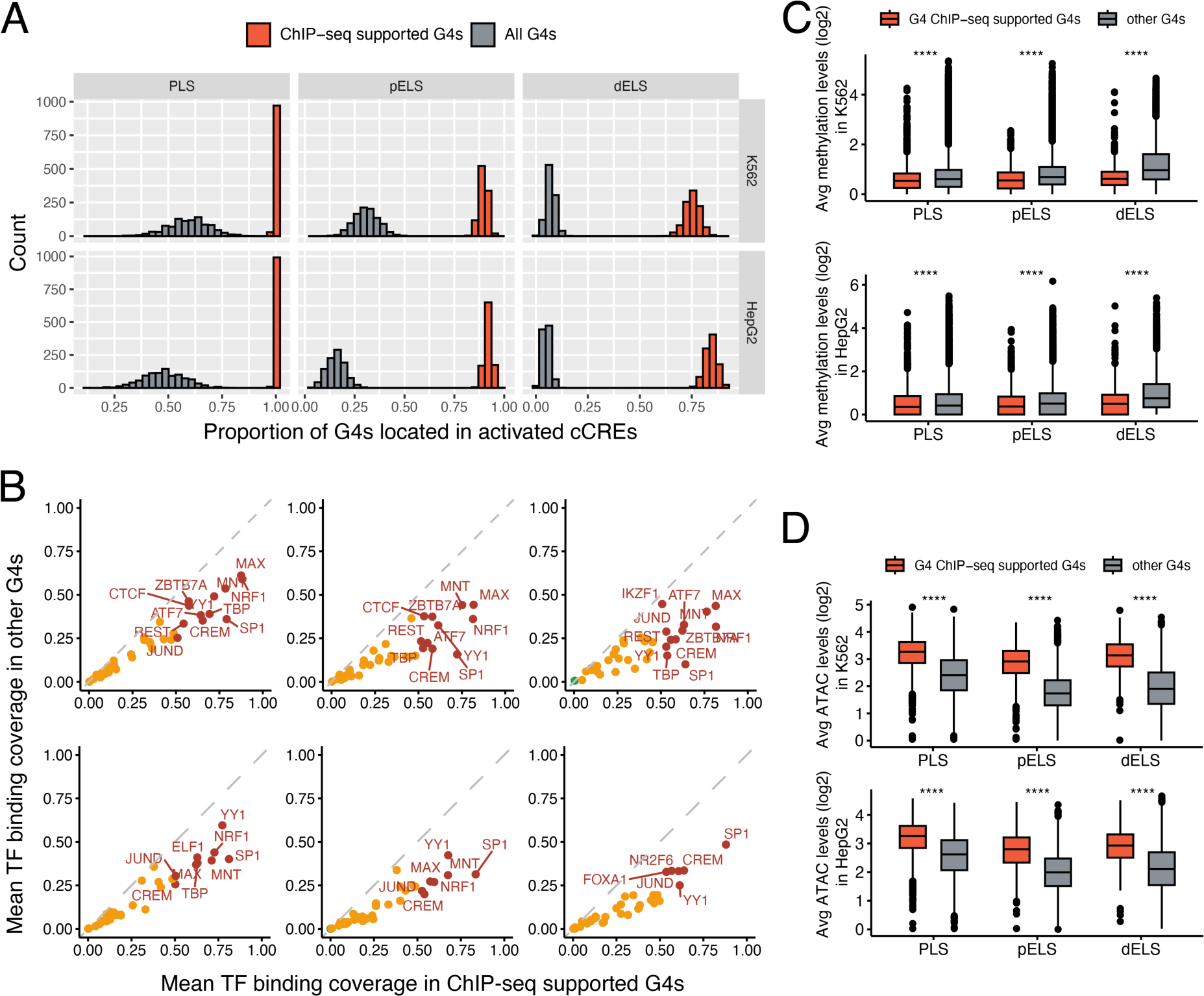
Comparing the association between G4 structures and G4 sequences with cCRE activation. (**A**) The histogram shows the statistics of the proportion of G4s located in the activated cCRE in each of the 1000 randomized experiments. Both G4 ChIP-seq supported G4s (orange) and background (all G4s; grey) proportions were estimated separately. Up-panel: K562 cell line; bottom-panel: HepG2 cell line. (**B**) Comparison of the average binding coverage of transcription factors on two types of G4s with and without G4 ChIP-seq experimental support. Red dots: the average binding coverage of this transcription factor on G4s with G4 ChIP-seq support is higher than that on other G4s and is higher than 0.5. Orange dots: similar to red dots, except that the average binding coverage is lower than 0.5. Green dots: the average coverage of this transcription factor binding on G4s with G4 ChIP-seq support is lower than that on other G4s. Top panel: the K562 cell line; bottom panel, the HepG2 cell line. (**C**) Differences in average methylation levels between K562 cell line (top panel) and HepG2 cell line (bottom panel) in regions (center ± 100bp) where endogenous G4s (with ChIP-seq support, orange) and other G4s (grey) are located. Wilcoxon test, ****: *P* < 0.0001. (**D**) Similar to (**C**) but calculating the differences in ATAC-seq signals.

To further investigate the connection between G4 structures, as opposed to sequences, and the activation of cCREs, we then performed a differential analysis of the transcription factor occupancy levels in the regions where two classes of G4s (with or without ChIP-seq support) are located. We found that almost all transcription factors preferentially bind to endogenous G4 regions (Fig. 6B), with some previously reported to interact with G4s, including SP1 [37, 38] and YY1 [39] (Fig. 6B). In addition, regions containing endogenous G4s on cCREs exhibit lower methylation levels (Fig. 6C; Wilcoxon test, *P* = 1.12×10^-34^, *P* = 1.19×10^-52^, *P* = 4.64×10^-112^ for comparisons of PLS, pELS, and dELS groups in K562 cells, respectively; *P* = 2.24×10^-8^, *P* = 6.02×10^-17^, *P* = 4.28×10^-41^ for comparisons of PLS, pELS, and dELS groups in HepG2 cells, respectively) and more open chromatin state (Fig. 6D; Wilcoxon test, *P* = 0, *P* = 0, *P* = 0 for comparisons of PLS, pELS, and dELS groups in K562 cells, respectively; *P* = 0, *P* = 6.71×10^-295^, *P* = 1.94×10^-179^ for comparisons of PLS, pELS, and dELS groups in HepG2 cells, respectively). We speculate that the G4 structures may regulate the activation of cCREs by shaping the local epigenetic environment.

### G4-associated cCREs are extensively activated in cancers

We then tested whether G4-associated cCREs would also be more inclined to be activated in cancer cells. To investigate this, we collected the pan-cancer chromatin accessibility profiles, along with the pan-cancer enhancer data inferred from the H3K27ac signals and the promoter activity estimated from the pan-cancer RNA-seq data (see methods).

We first examined the differences in the distribution density and the average coverage of ATAC-seq peaks between the two subgroups of cCREs, those associated with G4s or not, across various types of cancers. Our findings revealed that the distribution density as well as the average coverage of ATAC-seq peaks in the subgroup of G4-associated cCREs were higher than that of other cCREs in all cancer types (Fig. 7A-B, Supplementary Fig. S13-16). The proportion of G4-associated cCREs that overlapped with ATAC-seq peaks was also greater than that of the remaining cCREs in all types of cancers (Supplementary Fig. S17). This suggests that the presence of G4s in cCREs potentially correlates with a more accessible chromatin in cancers, thereby establishing a favorable environment for the activation of cCREs. Next, we evaluated the activation status of enhancer-like cCREs in various types of cancers. We discovered an elevated density of typical enhancers around G4-associated pELS elements in different cancers (Fig. 7C), while the disparity of enrichment is particularly prominent in the vicinity of dELS (Fig. 7D). Since the PLS element is potentially a *cis*-regulatory element acting as a promoter, we then assign PLS elements to their nearest promoters and directly substitute the PLS activity with promoter activity in various cancer tissues (see methods). As a result, G4-associated PLS elements exhibited relatively higher promoter activity (Fig. 7E; Wilcoxon test, *P* < 2.2×10^-16^ for all cancer types), independent of the type of cancer. Besides, a higher proportion of G4-associated PLS elements were highly activated across different cancer types (Supplementary Fig. S18).

**Figure 7.**
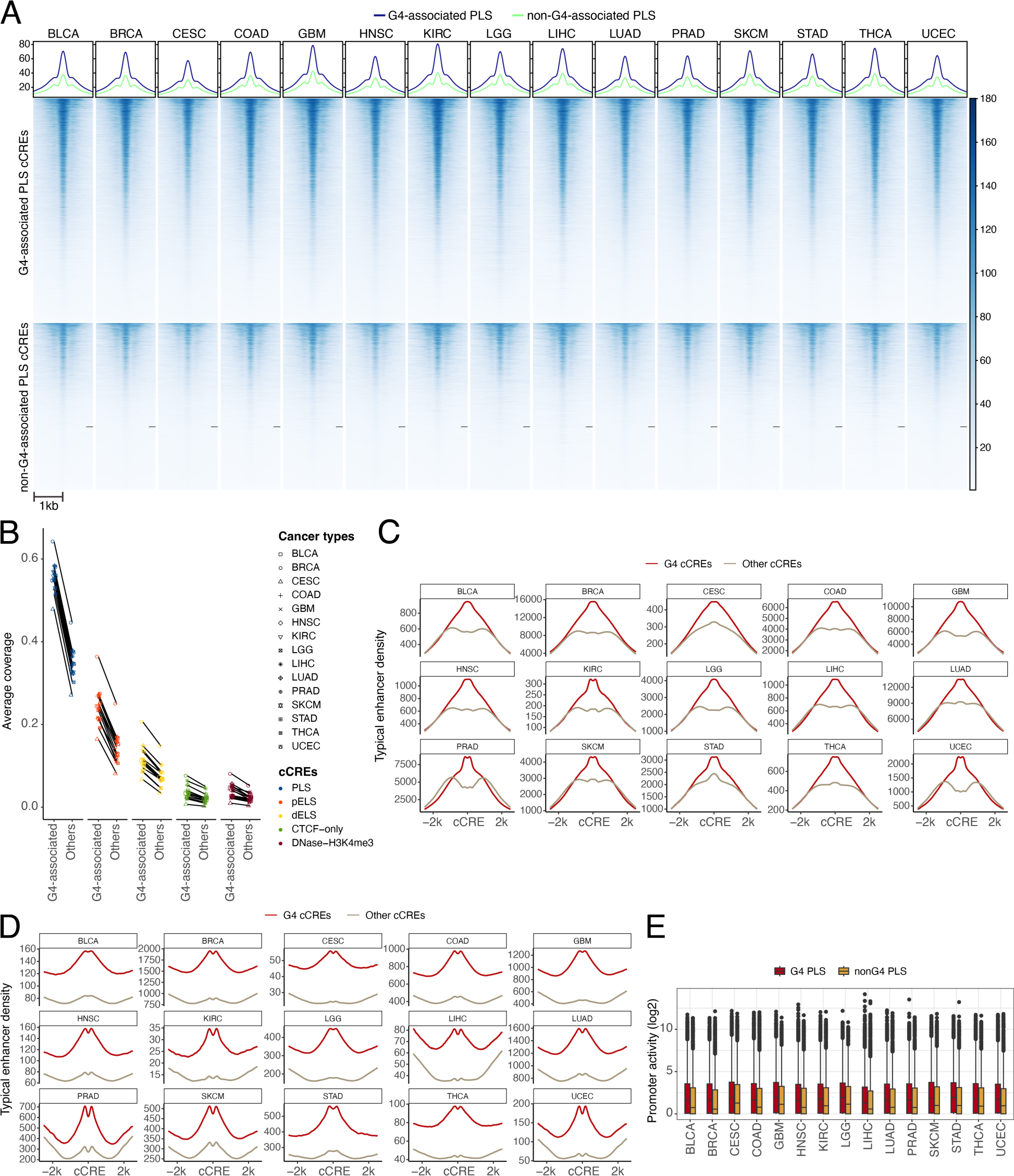
Quantification of the differences in pan-cancer chromatin accessibility, enhancer density, and promoter activity between G4-associated cCREs and other cCREs. **(A)** Heatmap and density plots of the ATAC-seq signals around G4-associated PLS elements and other (non-G4-associated) PLS elements. Heatmap legend indicating the ATAC-seq signal values around G4-associated PLS elements and non-G4-associated PLS elements in different cancer types. **(B)** The average coverage of pan-cancer ATAC-seq peaks in G4-associated cCREs and non-G4-associated cCREs. **(C)** The distribution of pan-cancer enhancers upstream and downstream of G4-associated pELS elements and non-G4-associated pELS elements is depicted. The Y-axis represents the magnitude of relative density. **(D)** Same as **(C)**, but the distribution of enhancers around dELS was examined. **(E)** Boxplot illustrates the disparities in promoter activity between G4-associated PLS elements (in red) and non-G4-associated PLS elements (in yellow) in different cancer types.

In conclusion, our analysis suggests that G4-associated cCREs are also susceptible to be activated and used in cancers.

## Discussion

Over the past decades, researchers have conducted extensive investigations into the physicochemical properties of G4s. However, in contrast, the study of the biological roles of G4s has only flourished in the last decade, thanks to the explosion of omics data. G4s are highly likely to become widely recognized as regulatory elements in the future. While relatively short, a G4 motif can mediate and regulate various biological functions in a non-coding manner due to its unique secondary structure. Our study addressed a fundamental question: are *cis*-regulatory elements in non-coding regions of the human genome be associated with G4s? In accordance with our research findings, we suggest that G4s are crucial components of some, but not all *cis*-regulatory elements in the human genome, and G4s may even constitute a core part of them, although not all cCREs are necessarily linked to G4s.

We noticed that the association between cCREs and G4s varies among different groups. Specifically, cCREs closer to TSS show a potentially stronger association with G4s, primarily including PLS and pELS elements. This is not unexpected, as G4s are inherently located near transcription start sites (TSSs), especially in upstream regions [40]. Interestingly, we observed that the G4Hunter scores of PLS-G4s tend to be the highest. G4Hunter scores can be used to characterize the stability of G4 structures, at least *in vitro*, with higher G4Hunter scores often indicating greater stability of G4 secondary structures [8]. Typically, G4s exert their biological functions mainly through the formation of folded secondary structures [41]. This aligns with their enrichment near TSSs, both suggesting that G4s on cCREs near TSSs potentially have stronger biological functions. Besides, these G4s are more closely associated with cCREs, possibly serving as their core functional elements, which is consistent with the results we obtained from a conservation perspective in mammalian genomes, namely that G4 G-runs near the TSS regions are more conserved compared to cCRE G-runs in the vicinity of TSSs. Strikingly, we only observe a weak association between CTCF-only cCREs and G4s, which was unexpected, as both our prior work and that of others had indicated a significant correlation between CTCF binding sites and G4 localization [27, 28]. Upon closer examination, it becomes evident that there are prerequisites for the association between G4s and CTCF-bound cCREs. When only CTCF signals are present, there is a weak association between the cCREs and G4s; however, when the CTCF-bound cCREs also possess H3K4me3 or H3K27ac signals, a stronger association with G4s emerges. This strong association cannot be simply explained as a correlation between G4s and H3K4me3 or H3K27ac signals themselves, because when we classified cCREs such as PLS based on CTCF alone, a higher proportion of cCREs containing CTCF signals would be associated with G4s, especially in PLS and pELS elements. This difference becomes even more pronounced when we directly observe cCREs and G4s from the K562 and HepG2 cell lines. We speculate that the interaction between CTCF and G4 in regions such as PLS or ELS may contribute to the formation of chromatin loops and other chromatin conformations, thereby facilitating the function of PLS or ELS elements.

Previous studies have suggested an association between G4s and various epigenetic marks [33, 34, 42]. In this study, we showed that the presence of G4s is often linked to the local chromatin environment that supports the activation or functioning of cCREs, such as increased occupancy of transcription factors and a higher unmethylated state. This prompts us to formulate a hypothesis that some cCREs may become regulatory elements precisely because of the presence of G4s. In other words, the presence of G4s is an important factor ensuring their biological functions. Intriguingly, we found that it is the G4 structures rather than the sequences that is associated with the activation of cCREs. The G4 structure regions often display higher transcription factor occupancy as well as lower methylation levels, creating a favorable epigenetic environment for the activation of cCREs. Nevertheless, the relationship between the formation of G4 secondary structures and changes in the epigenetic environment is akin to a ‘chicken and egg’ problem, rendering it difficult for us to make causal inferences *via* computational methods. Yet, existing research has provided us with some clues. For instance, it has been confirmed that G4 structures can shape the local hypomethylation landscape of the genome [43]. Additionally, G4 structures can interact transcription factors, serving as hubs for their bindings [36]. Investigating how G4 structures influence the local chromatin environment and, consequently, impact *cis*-regulatory elements will be a very intriguing topic.

Interpreting the consequences of variants in noncoding regions has long been a challenging endeavor [44, 45]. Our study indicates that variants on G4s, especially those on G-runs that are likely to be important for the formation of G4 structures, tend to possess a higher regulatory potential and tend to be conserved, as substitutions often lead to deleterious variants. This indirectly underscores the importance of G4s in cCREs. Thus, we recommend that G4 should be considered as a feature when developing new models or algorithms to predict the regulatory capacity of non-coding regions.

We also noted that in multiple cancers, G4-associated cCREs are indeed more active, as evidenced by higher levels of chromatin accessibility, promoter activity, as well as enhancer activity. In view of the fact that *cis*-regulatory elements also regulate the transcriptome expression of cancer cells or tissues, it would be crucial in the future to further identify those G4-associated cCREs that are aberrantly activated compared to adjacent non-cancerous tissues. We also look forward to the extensive application of G4 ChIP-seq technology in the identification of G4 secondary structures in cancer tissues as well as their paired normal tissues. While identifying G4s in some normal cell lines that match the cancer cell line type is essential. This will enable us to understand which G4 structures are abnormally formed in cancer, whether these abnormally formed G4 structures can affect their associated *cis*-regulatory elements, and ultimately, whether they can influence the regulation of target genes, potentially leading to tissue carcinogenesis. However, due to the limitations in G4 sequencing technology, we are currently unable to obtain such sample data and analyze the structural alteration of G4s in cancer tissues in conjunction with the activation status of cCREs.

In conclusion, our study suggests that G4s are crucial regulatory components of cCREs. Our finding lays the groundwork for a deeper comprehension of *cis*-regulatory elements and the exploration of their underlying mechanisms in future research studies.

## Data availability

The human genome G4 sequences predicted by G4Hunter in this study have been deposited at https://github.com/rongxinzh/G4_CRE/raw/main/G4Hunter_w25_s1.5_hg38.txt. The annotation of the presence of G4s on cCREs can be accessed through https://github.com/rongxinzh/G4_CRE/raw/main/G4_cCRE_annotation.txt.

## Code availability

Source code for this project is available at https://github.com/rongxinzh/G4_CRE.

## Abbreviations

Abbreviation: Meaning
G4: G-quadruplex
cCRE: candidate *cis*-regulatory element
PLS: promoter-like signatures
pELS: proximal enhancer-like signatures
dELS: distal enhancer-like signatures
CTCF-only: high DNase and CTCF signals only
DNase–H3K4me3: high DNase and H3K4me3 signals only
BLCA: Bladder Urothelial Carcinoma
BRCA: Breast Invasive Carcinoma
CESC: Cervical Squamous Cell Carcinoma
COAD: Colon Adenocarcinoma
GBM: Glioblastoma Multiforme
HNSC: Head and Neck Squamous Cell Carcinoma
KIRC: Kidney Renal Clear Cell Carcinoma
LGG: Low Grade Glioma
LIHC: Liver Hepatocellular Carcinoma
LUAD: Lung Adenocarcinoma
PRAD: Prostate Adenocarcinoma
SKCM: Skin Cutaneous Melanoma
STAD: Stomach Adenocarcinoma
THCA: Thyroid Carcinoma
UCEC: Uterine Corpus Endometrial Carcinoma

