## Supplementary Materials for "G-quadruplexes as pivotal components of *cis*-regulatory elements in the human genome"

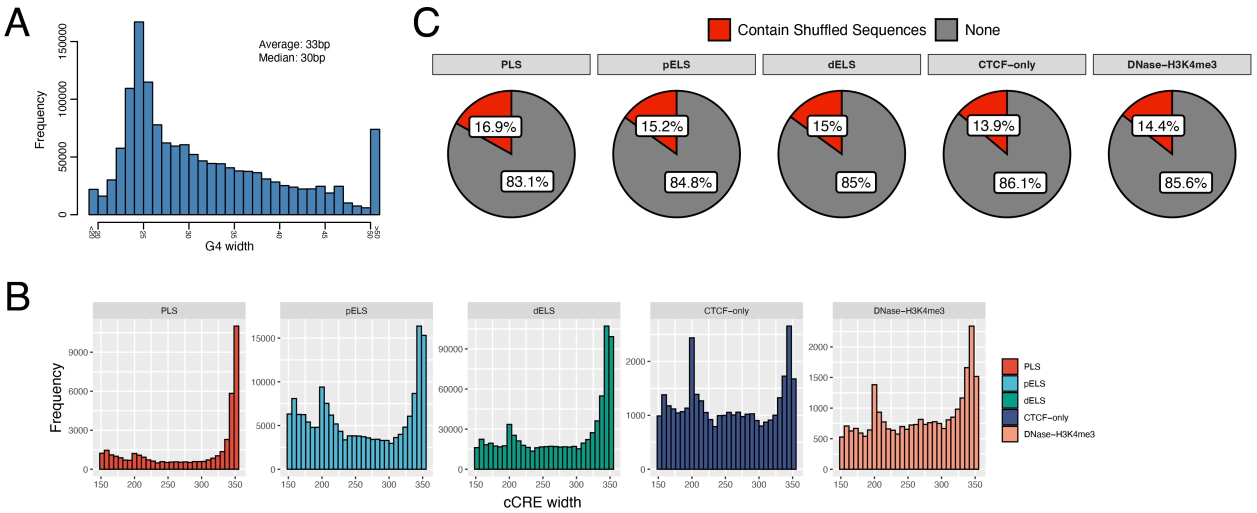


Supplementary Figure S1. (**A**) The widths of G4 sequences in the human genome predicted by the G4Hunter software. The x-axis displays the width of G4s, and the y-axis shows the frequency of G4 sequences with that width. The leftmost and rightmost bars represent the number of G4s with widths less than 20 and greater than 50, respectively. (**B**) Frequency statistics of the widths of the five cCREs groups. The x-axis represents the width of cCREs and the y-axis represents the frequency of that width. (**C**) Pie charts shows the proportion of cCREs that contain shuffled sequences as well as those that do not.

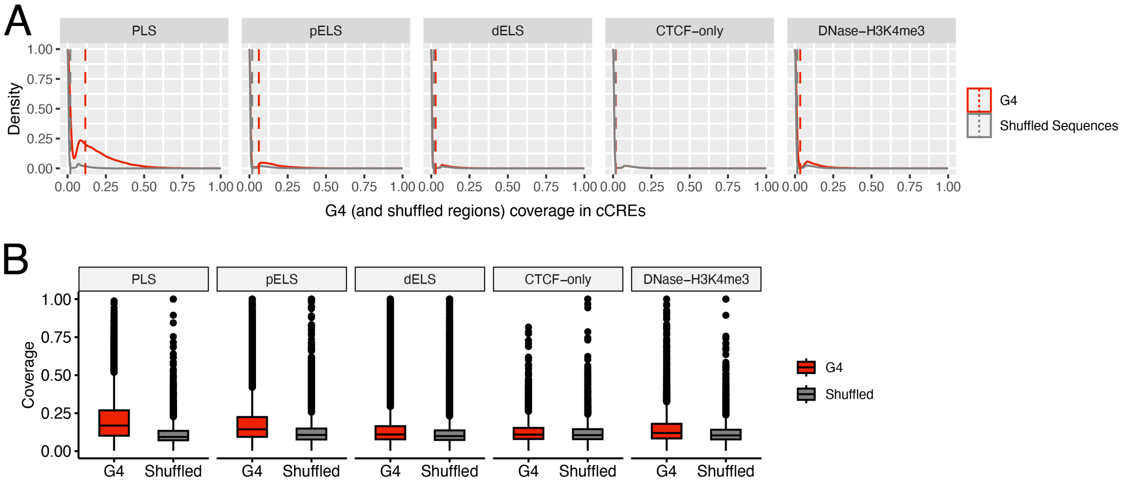


Supplementary Figure S2. Comparison of G4 and shuffled sequence coverage on cCREs. (**A**) Density plot shows the distribution of the coverage of G4s or shuffled sequences on all cCREs of different groups. The red and the grey dashed lines represent the average coverage of G4s on the cCREs as well as the average coverage of shuffled sequences, respectively. (**B**) Comparison of the G4 coverage in G4-contained cCREs with the shuffled sequence coverage in shuffled-sequence-contained cCREs across five different cCRE groups.

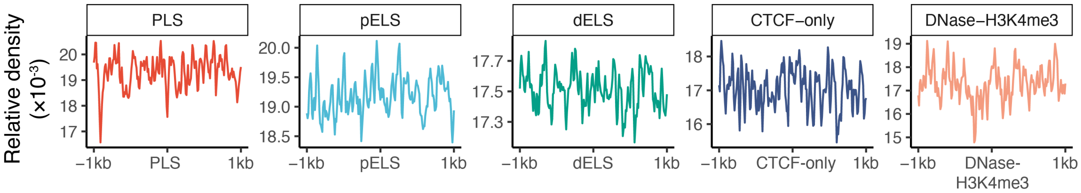


Supplementary Figure S3. Relative density of shuffled sequences around (±1kb) five groups of cCREs.

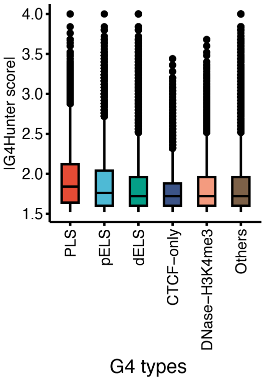


Supplementary Figure S4. The absolute G4Hunter scores for G4s located in different groups of cCREs. The group of others represents those G4s that cannot be annotated into the five groups of cCREs.

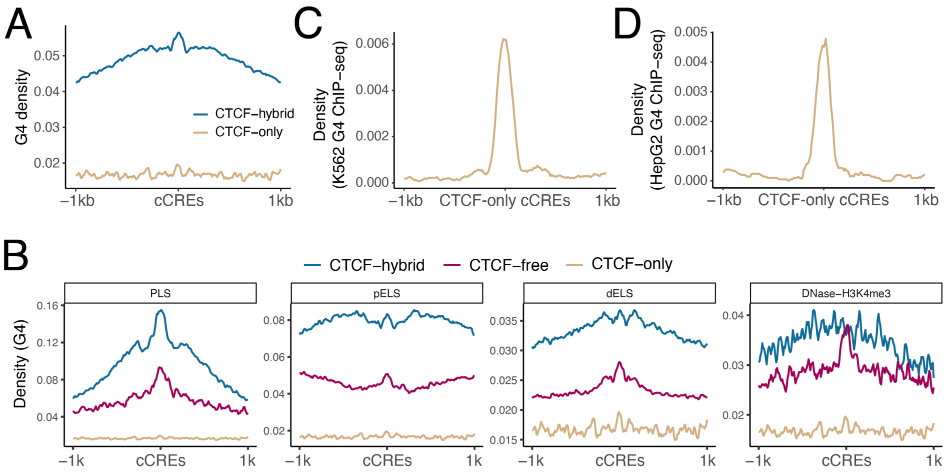


Supplementary Figure S5. Distribution of G4s in cCREs with different CTCF occupancy patterns. (**A**) Relative density of G4s (predicted by G4Hunter software) around (±1kb) CTCF-hybrid cCREs as well as CTCF-only cCREs. The blue and brown curves depict the distribution of G4s around CTCF-hybrid and CTCF-only cCREs, respectively. (**B**) Relative density of G4s (predicted by G4Hunter software) around (±1kb) CTCF-hybrid cCREs and CTCF-free cCREs. CTCF-only cCREs were also included for comparison. (**C-D**) Relative density of G4 ChIP-seq high confidence peaks around CTCF-only cCREs in the K562 (**C**) and HepG2 (**D**) cell lines.

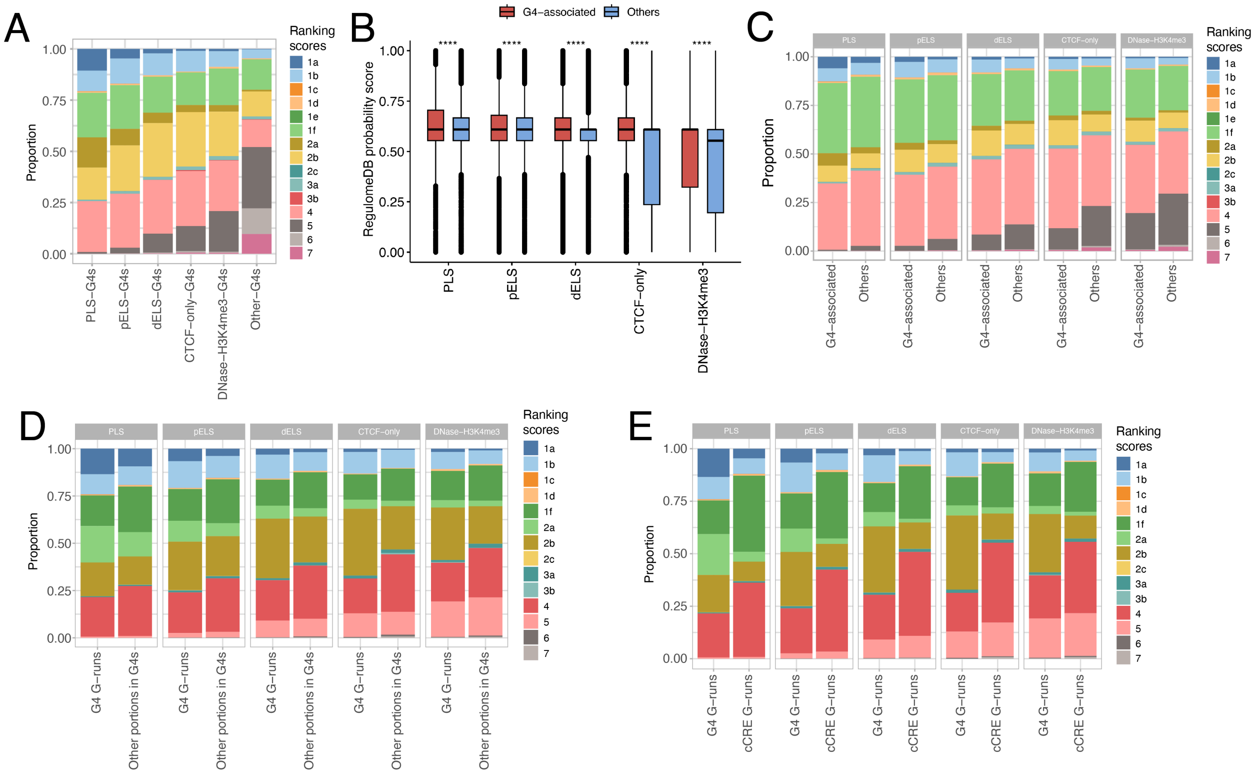


Supplementary Figure S6. Overview of RegulomeDB ranking and probability scores in G4s and cCREs. (**A**) The proportion of variants with different RegulomeDB ranking scores in different groups of G4s. (**B**) The differences of RegulomeDB probability scores for variants located in G4-associated cCREs and other (non-G4-associated) cCREs. Wilcoxon test, ****: $P < 0.0001$ (**C-E**) Comparison of the proportion of variants with different RegulomeDB ranking scores between G4-associated and other cCREs (**C**). (**D**) and (**E**) are similar to (**C**), with the main differences being that in (**D**), the comparisons are performed between G4 G-runs and other portions in G4s, while in (**E**), the comparisons are made between G4 G-runs and cCRE G-runs.

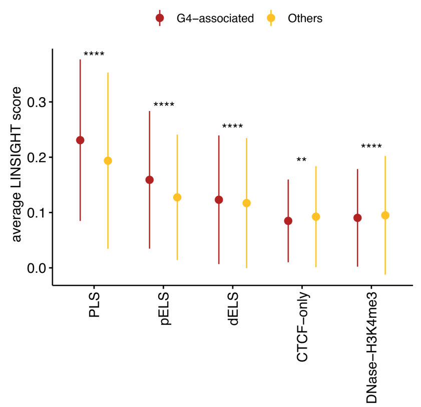


Supplementary Figure S7. The differences in average LINSIGHT scores between G4-associated cCREs and other (non-G4-associated) cCREs. Wilcoxon test, **: *P* < 0.01, ****: *P* < 0.0001.

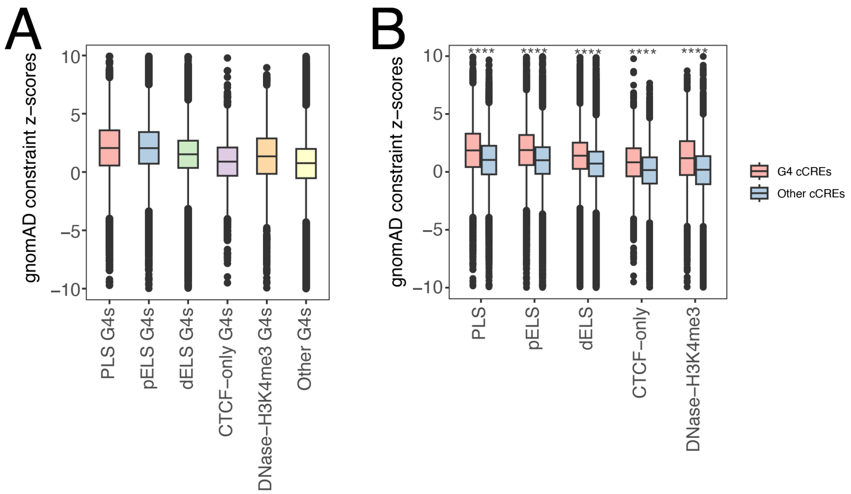


Supplementary Figure S8. (**A**) The differences in gnomAD constraint z-scores for G4s located in different cCREs. (**B**) Boxplot shows the disparities in gnomAD constraint z-scores between G4-associated cCREs and non-G4-associated cCREs. Wilcoxon test, ****: *P* < 0.0001.

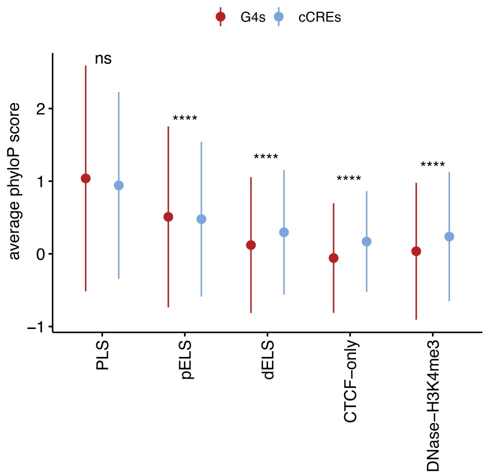


Supplementary Figure S9. Error plots show the differences in average phyloP scores between cCREs and the G4s overlapped with the corresponding cCREs. Dots and ranges denote mean and standard deviation, respectively. Wilcoxon test, ns: $P > 0.05$, ****: $P < 0.0001$.

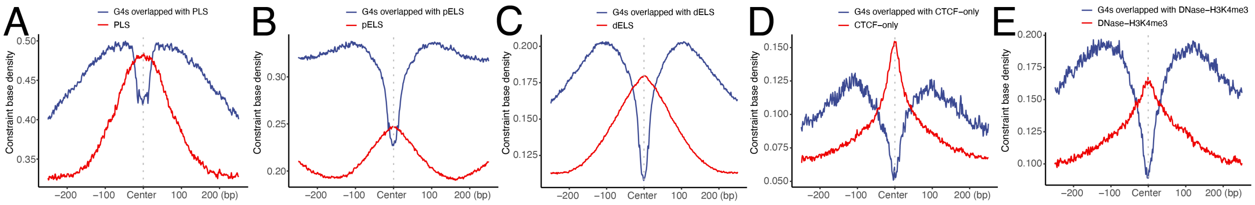


Supplementary Figure S10. (**A-E**) The distribution of constraint bases around different groups of cCREs (red curve) as well as around the G4s that overlapped with the corresponding group of cCREs (blue curve).

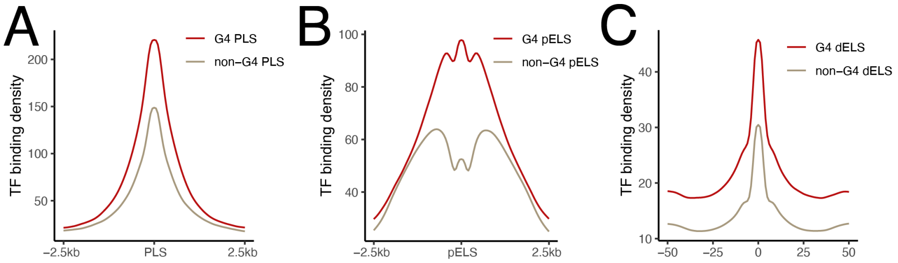


Supplementary Figure S11. TF binding density around G4-associated (red curve) and non-G4-associated (brown curve) PLS, pELS, and dELS cCREs.

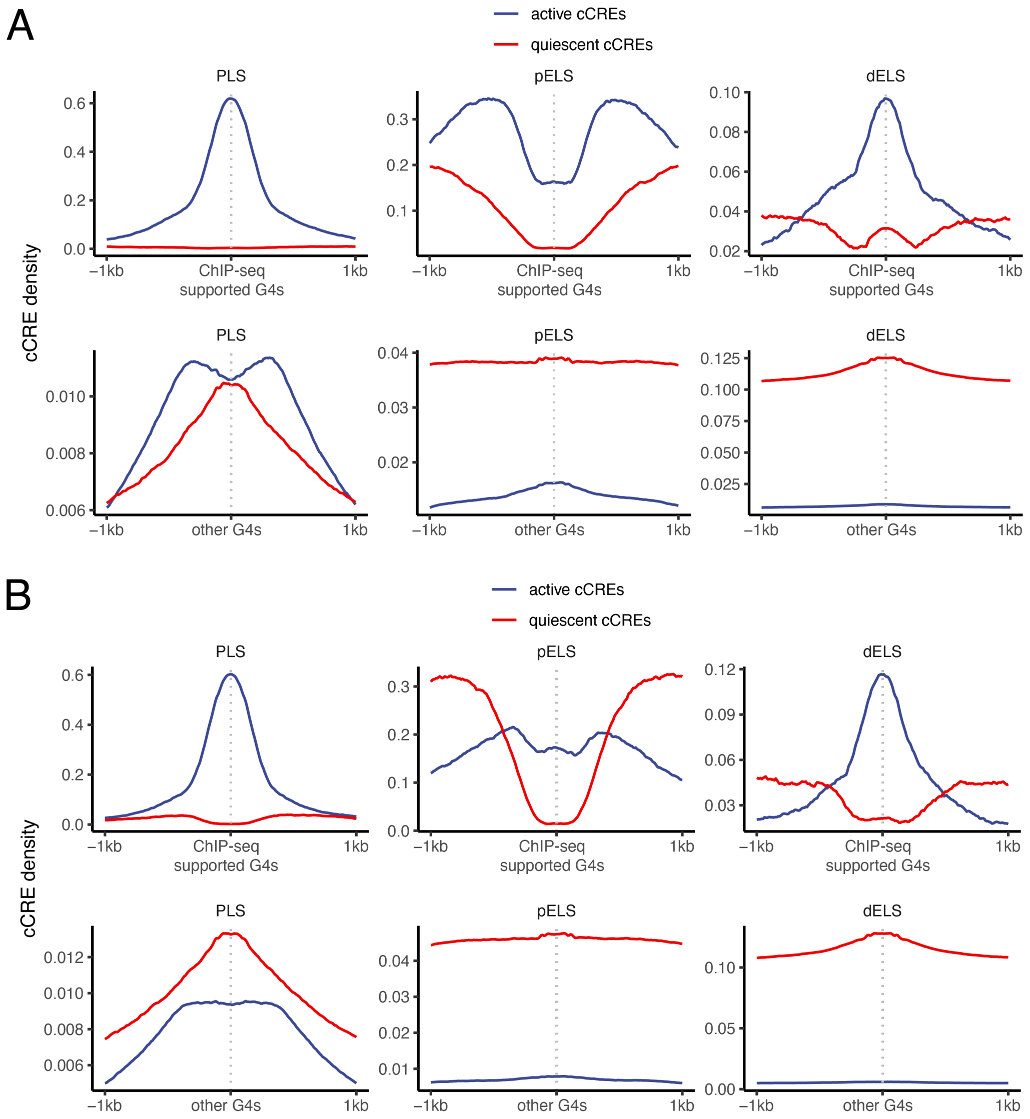


Supplementary Figure S12. (**A**) Distribution of activated cCRE (red curves) and quiescent cCRE (blue curves) around (±1kb) G4s supported by G4 ChIP-seq experiments, as well as other G4s in the K562 cell line. (**B**) Similar to (**A**), but in the HepG2 cell line.

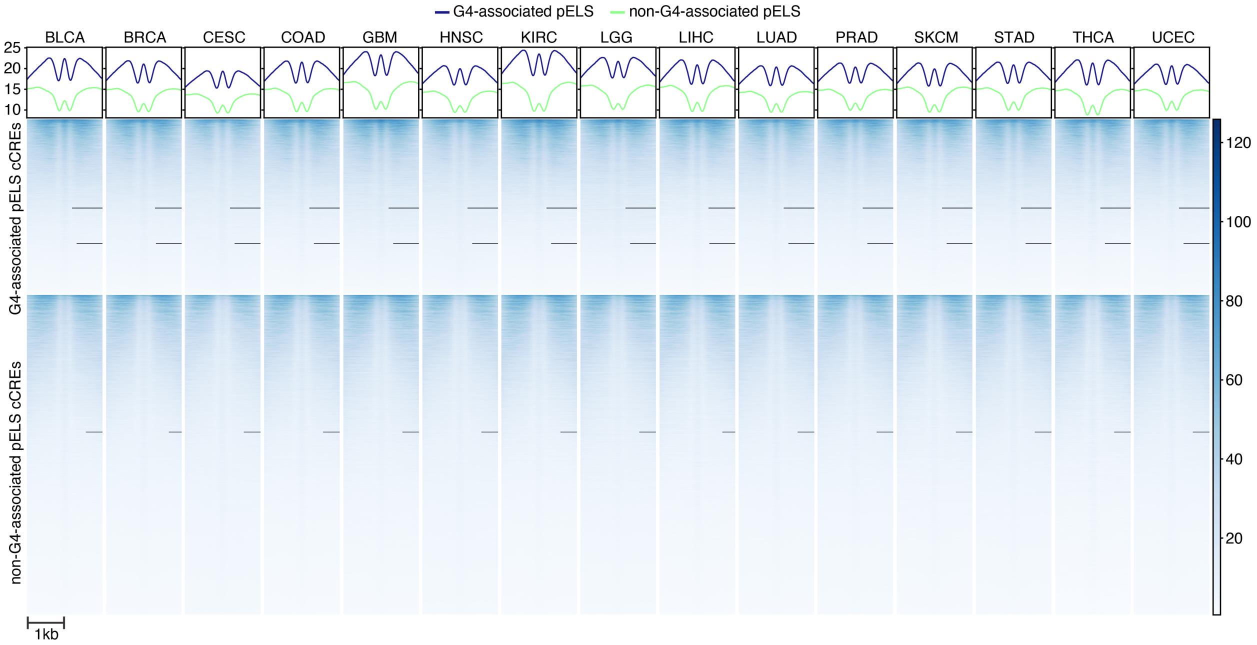


Supplementary Figure S13. Heatmap shows the pan-cancer ATAC-seq signals around G4-associated pELS elements and non-G4-associated pELS elements. Heatmap legend illustrates the signaling levels of ATAC-seq.

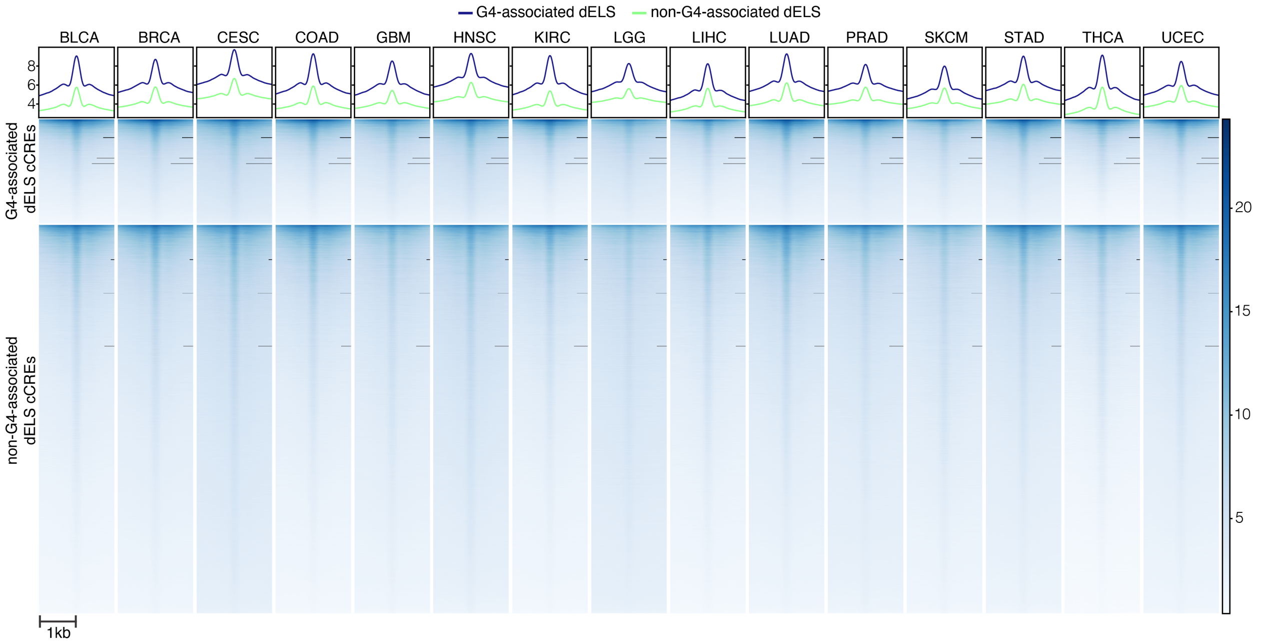


Supplementary Figure S14. Heatmap shows the pan-cancer ATAC-seq signals around G4-associated dELS elements and non-G4-associated dELS elements. Heatmap legend illustrates the signaling levels of ATAC-seq.

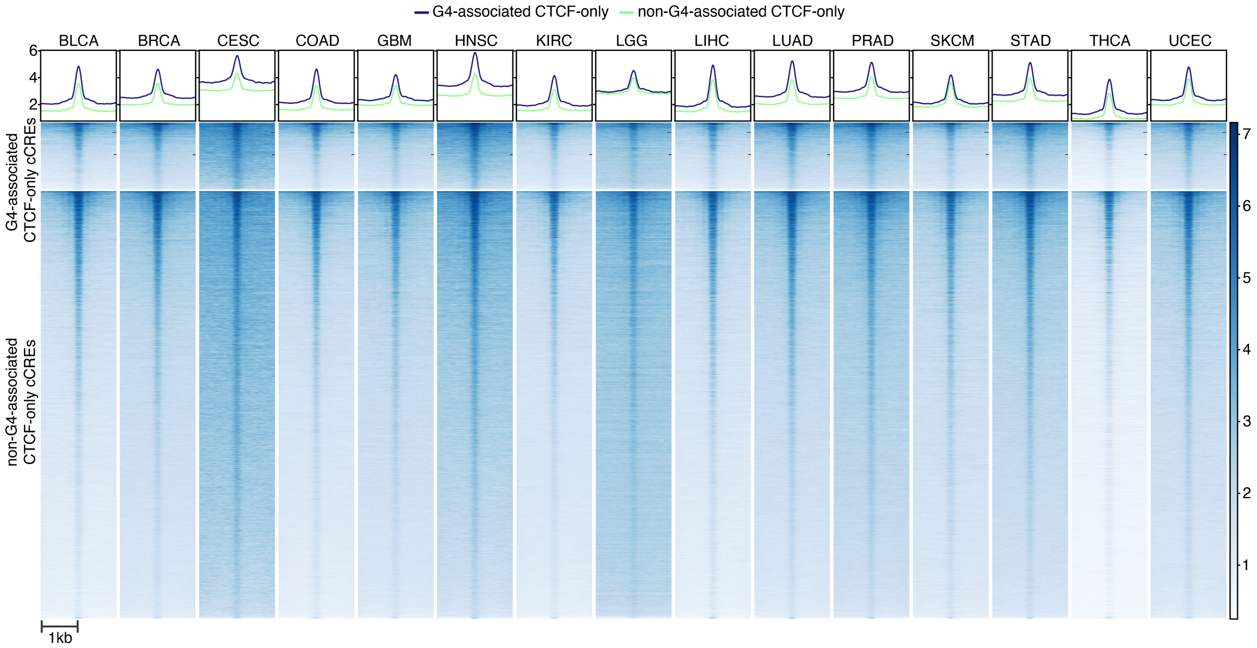


Supplementary Figure S15. Heatmap shows the pan-cancer ATAC-seq signals around G4-associated CTCF-only elements and non-G4-associated CTCF-only elements. Heatmap legend illustrates the signaling levels of ATAC-seq.

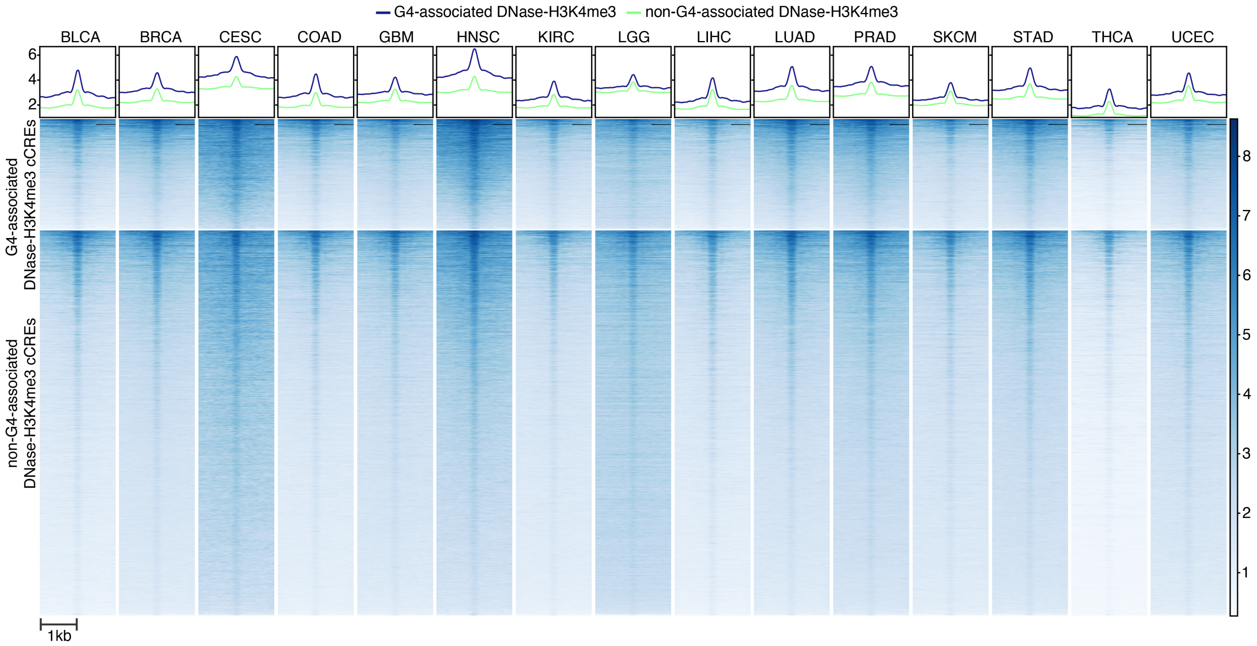


Supplementary Figure S16. Heatmap shows the pan-cancer ATAC-seq signals around G4-associated DNase-H3K4me3 elements and non-G4-associated DNase-H3K4me3 elements. Heatmap legend illustrates the signaling levels of ATAC-seq.

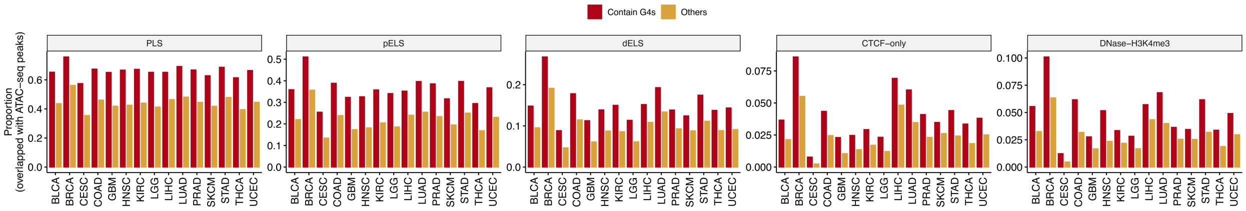


Supplementary Figure S17. Bar plot indicates the proportion of G4-associated cCREs (red) and non-G4-associated cCREs (yellow) that overlapped with ATAC-seq peaks in distinct cancer types.

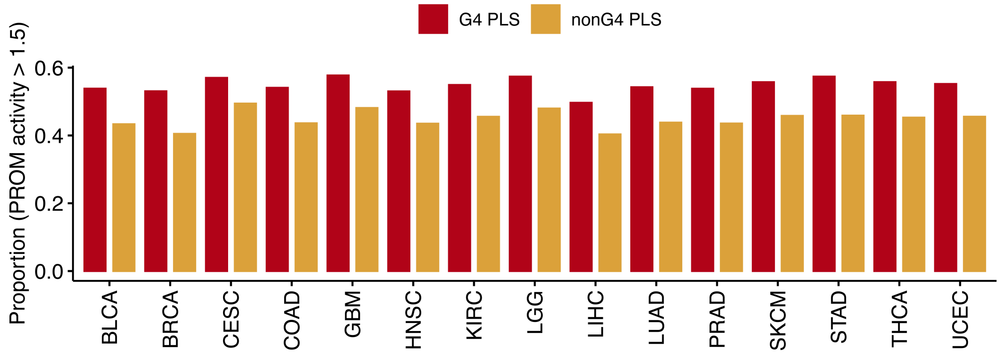


Supplementary Figure S18. Proportion of G4-associated PLS (red) and non-G4-associated PLS (yellow) with activity greater than 1.5 (highly activated) in different cancer types.
